# Supervised learning of high-confidence phenotypic subpopulations from single-cell data

**DOI:** 10.1101/2023.03.23.533712

**Authors:** Tao Ren, Canping Chen, Alexey V. Danilov, Susan Liu, Xiangnan Guan, Shunyi Du, Xiwei Wu, Mara H. Sherman, Paul T. Spellman, Lisa M. Coussens, Andrew C. Adey, Gordon B. Mills, Ling-Yun Wu, Zheng Xia

**Author notes:** Corresponding authors: Zheng Xia, Ph.D.; Ling-Yun Wu, Ph.D.

## Abstract

Accurately identifying phenotype-relevant cell subsets from heterogeneous cell populations is crucial for delineating the underlying mechanisms driving biological or clinical phenotypes. Here, by deploying a learning with rejection strategy, we developed a novel supervised learning framework called PENCIL to identify subpopulations associated with categorical or continuous phenotypes from single-cell data. By embedding a feature selection function into this flexible framework, for the first time, we were able to select informative features and identify cell subpopulations simultaneously, which enables the accurate identification of phenotypic subpopulations otherwise missed by methods incapable of concurrent gene selection. Furthermore, the regression mode of PENCIL presents a novel ability for supervised phenotypic trajectory learning of subpopulations from single-cell data. We conducted comprehensive simulations to evaluate PENCIL’s versatility in simultaneous gene selection, subpopulation identification and phenotypic trajectory prediction. PENCIL is fast and scalable to analyze 1 million cells within 1 hour. Using the classification mode, PENCIL detected T-cell subpopulations associated with melanoma immunotherapy outcomes. Moreover, when applied to scRNA-seq of a mantle cell lymphoma patient with drug treatment across multiple time points, the regression mode of PENCIL revealed a transcriptional treatment response trajectory. Collectively, our work introduces a scalable and flexible infrastructure to accurately identify phenotype-associated subpopulations from single-cell data.

## Introduction

Heterogeneous cellular systems alter cell states and compositions in response to development, perturbations, pathological change, and clinical intervention, resulting in phenotypically distinct cell subpopulations^1–4^. Rapidly accumulating single-cell studies are profiling samples from different experimental or pathological conditions, such as wild-type vs. knockout conditions^5^, treatment resistance vs. responder groups^6^, disease progression graded with scores^7^, and treatment across multiple time points^8^. Distinguishing subpopulations associated with phenotypes of interest from heterogeneous cell populations will improve phenotype-specific gene signal detection and enables reliable downstream interrogation of phenotypic cell types and states, which is a key step in delivering knowledge from the designed single-cell experiments. Therefore, it is essential to develop analytical tools to identify phenotypic subpopulations from single-cell data.

For categorical phenotypes, the phenotype-associated subpopulations can be identified through differential abundance analysis. A straightforward method is to cluster cells first and then compare the ratios of conditions in each cluster^9^. Such clustering-based methods, however, depend on the subjective clustering step and are often suboptimal because the phenotype-specific subpopulations are usually not detected by standard clustering methods. Therefore, recent developments have proposed clustering-free strategies like DAseq^10^, Milo^11^, and MELD^12^ by examining phenotype labels of cells connected through the k-nearest neighbor (KNN) graph. Nevertheless, KNN graphs require gene selection beforehand, which are determined separately in an unsupervised manner, e.g., the top most variable genes.

Such unsupervised gene selection approaches^13, 14^ may not capture phenotype-associated cell subpopulations hidden in a latent gene space. As a result, to accurately detect the cells of interest, gene selection must be embedded into the subpopulation identification process. However, given the cell-cell similarity matrix as input, the KNN-based tools cannot incorporate gene selection into subpopulation identification, leaving the two integral steps separated.

Moreover, beyond detecting static categorical cell subsets, we need to order the selected cells along the continuous phenotypic trajectory to reveal transitions and relationships during dynamic biological processes, such as tissue development and disease progression^15–20^, a critical task for single-cell analysis^21^. However, although Milo^11^ can input continuous phenotypes, it only interprets subpopulations increasing or decreasing with the phenotype qualitatively without ordering cells in a trajectory manner. As a result, further methodological development of new frameworks beyond cell-cell similarity is necessary.

In order to select informative genes, we need a framework that can directly take the gene matrix as an input. Additionally, this new framework must reject irrelevant cells while retaining high-confidence cells. To address these two needs, we propose a new tool that uses the learning with rejection (LWR) strategy to detect high-confidence **<u>p</u>**h**<u>en</u>**otype-asso**<u>c</u>**iated subpopulat**i**ons from single-cel**l** data (PENCIL). LWR includes a prediction function (Fig. 1a) along with a rejection function (Fig. 1b) to reject low-confidence cells.

**Fig. 1.**
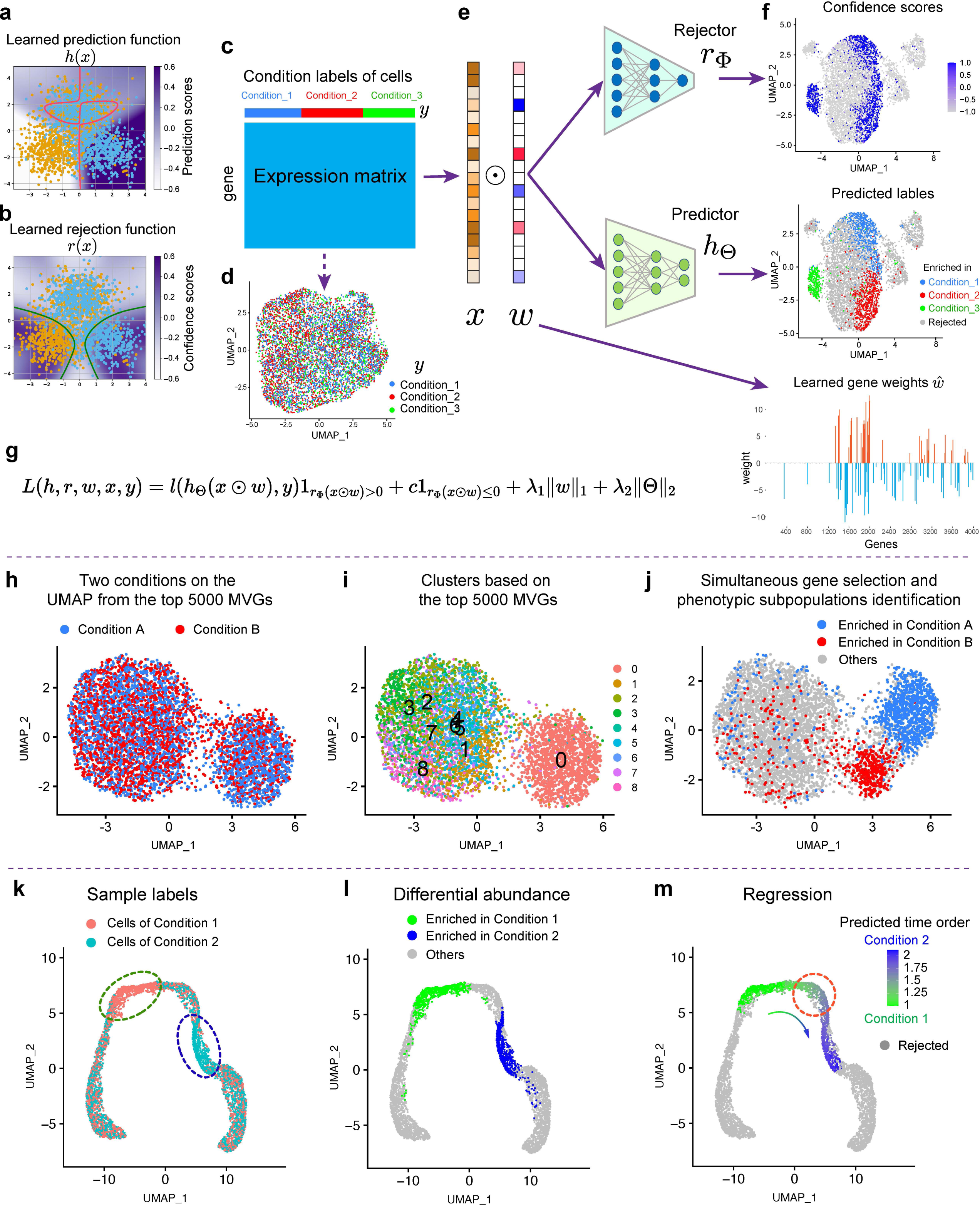
The workflow of PENCIL and its main functions. **a-b**, A simulated example to show the learned prediction model with the red line as the boundary with prediction scores ℎ(*x*) = 0 to separate the two predicted classes; and the learned rejection model with the green lines as the boundary with confidence scores *r*(*x*) = 0 to reject cells. **c**, The inputs for PENCIL are a single-cell data matrix and condition labels of all cells *y*. **d**, The single-cell expression matrix is visualized by the UMAP using the top 2000 most variable genes (MVGs) with cells colored by the condition labels. **e,** The three trainable components of PENCIL: gene weights *w*, rejector module, and predictor module. **f,** The outputs of PENCIL are confidence scores, predicted labels, and learned gene weights. The UMAPs are generated by the PENCIL selected genes with *ŵ* ≠ 0. **g,** The rejection-based total loss function of PENCIL for the optimization. **h,** UMAP using the top 5000 MVGs showing a dataset with two conditions colored by their condition labels. **i,** Standard clustering analysis based on the top 5000 MVGs. **j,** UMAP based on the PENCIL selected genes showing the identified phenotype-enriched cell subpopulations. **k,** UMAP visualization of a simulated single-cell RNA-seq data with cells colored by the conditions. The designated regions enriched in each condition were denoted by the dashed ovals. **l**, Differential abundance analysis like Milo and classification mode of PENCIL can only identify static phenotype-associated cell subpopulations from the data shown in **k**. **m,** Continuous phenotype regression PENCIL analysis rejected the irrelevant cells and predicted the time orders of phenotypic cells to reveal continuous transition states as indicated by the red dashed circle.

Then, by embedding a feature selection function into this LWR framework, PENCIL can perform gene selection during the training process, which allows learning proper gene spaces that facilitate accurate subpopulation identifications from single-cell data. Thus, the PENCIL framework also provides a new perspective for gene selection in single-cell analysis beyond the unsupervised architecture. Furthermore, by updating the prediction loss function, PENCIL has the flexibility to address various phenotypes such as binary, multi-category and continuous phenotypes. Most importantly, the regression mode of PENCIL can order cells to reveal the subpopulations undergoing continuous transitions between conditions, which is fundamentally different from the differential abundance analysis. To our knowledge, PENCIL represents the first tool for simultaneous gene selection and phenotype-associated subpopulation identification from single-cell data that can detect subpopulations enriched by specific categorical phenotypes or learn their continuous phenotypic trajectory.

## Results

### Overview of PENCIL

To construct a new framework distinct from the existing KNN-based frameworks, we introduced a learning with rejection (LWR) strategy (Fig. 1a,b) into single-cell data analysis for phenotypic subpopulation identification. Then, by incorporating a feature selection function into LWR, we developed a new tool named PENCIL to simultaneously select genes and identify phenotype-associated cell subpopulations from single-cell data. The data sources for PENCIL input include a single-cell quantification matrix and condition labels for all cells (Fig. 1c,d). Condition labels can be in various forms, such as multiple experimental perturbations, disease stages, time points, and so on. In brief, PENCIL consists of three modules, gene weights, predictor, and rejector (Fig. 1e). Gene weights are penalized with a sparse penalty (*l*_1_-norm) to select genes informative for the targeted phenotypes; the predictor is a general trainable model in supervised learning that is used to predict cell labels, and the rejector assigns each cell with a confidence score to quantify the credibility of the predicted label from the predictor (Fig. 1f). The parameters of all three modules are trained by minimizing the total loss function and regularization terms on the input expression matrix with condition labels (Fig. 1g). Then, the combination of the predicted labels and the confidence scores (*r*(*x*) > 0) from the rejection function will output the selected subpopulations with predicted labels (Methods).

PENCIL is flexible to take either categorical phenotypes or continuous variables as inputs by changing the prediction function. For example, Figure 1h shows a simulated scRNA-seq dataset with binary phenotype labels in a Uniform Manifold Approximation and Projection (UMAP)^22^ using the top 5000 most variable genes (MVGs). The standard top 5000 MVGs based clustering analysis cannot distinguish the two phenotypic clusters contained in cluster 0 (Fig. 1i). In contrast, our classification mode of PENCIL with gene selection can identify the two subtle phenotypic subpopulations, as shown by the UMAP based on the PENCIL selected genes (Fig. 1j), demonstrating the importance of gene selection in cell subpopulation identification. Furthermore, by setting the predictor module as a regressor, PENCIL can handle continuous phenotype labels like time points and disease stages, which carries out a fundamentally different task than the differential abundance analysis in the classification mode for single-cell applications. For instance, in a simulated single-cell dataset from two conditions^23^ (Fig. 1k), the category-based subpopulation identification methods, like Milo^11^ or the classification mode of PENCIL, can only identify the differentially abundant subpopulations (Fig. 1l). Intriguingly, the regression-based PENCIL can reconstruct the phenotypic trajectory to reveal the subpopulations that are undergoing a continuous transition between conditions (Fig. 1m), like the cells transforming from normal to malignant. Thus, the regression mode of PENCIL offers an opportunity to understand dynamic processes of biology and disease that is unattainable with existing methods.

### PENCIL’s classification mode simultaneously selects genes and cells

To test the effectiveness of PENCIL, we set up a series of simulated datasets for the classification task, and performed comprehensive comparisons with existing methods, including DAseq^10^, Milo^11^, and MELD^12^. We exploited a real T cell scRNA-seq expression dataset^6^ with 6,350 cells to generate various simulation settings by picking informative gene sets and simulating condition labels accordingly. In the simulation with two conditions, we first selected a subset of genes from the top 2000 most variable genes (MVGs) as informative genes for the downstream clustering and visualization in UMAP to generate ground truth phenotypic subpopulations. After clustering based on these manually selected genes, we picked out two clusters and designated them to be ground truth subpopulations enriched in specific conditions, respectively (Fig. 2a), and all other cells were set as background cells. Next, we assigned condition labels to the cells based on the ground-truth subpopulations and background cells. For each ground-truth subpopulation, we used a number *α* called mixing rate to control the ratio between the majority and the minority condition labels. Within each ground truth subpopulation, we assigned (1 − *α*) of the total cells with the designated majority condition label, and the remaining cells with other labels. For the background cells, each cell was randomly assigned with a condition label. In this way, we generated the condition labels for all cells for one simulation, as shown in Figure 2b with a mixing rate *α* = 0.1 (see Supplementary Figure 1 and the Methods for more details). Since the genes to generate the clustering and UMAP are only a subset of the total genes, the standard scRNA-seq analysis pipeline using the top 2000 MVGs will not capture the proper cell similarities, resulting in ambiguous aggregation patterns for cell label information (Fig. 2c,d), thus making it difficult for the methods using the KNN based on the top 2000 MVGs to identify subpopulations of interest. After setting up the simulation, we used the gene expression matrix of the top 2,000 MVGs and the simulated conditions labels as the source data for all four methods.

**Fig. 2.**
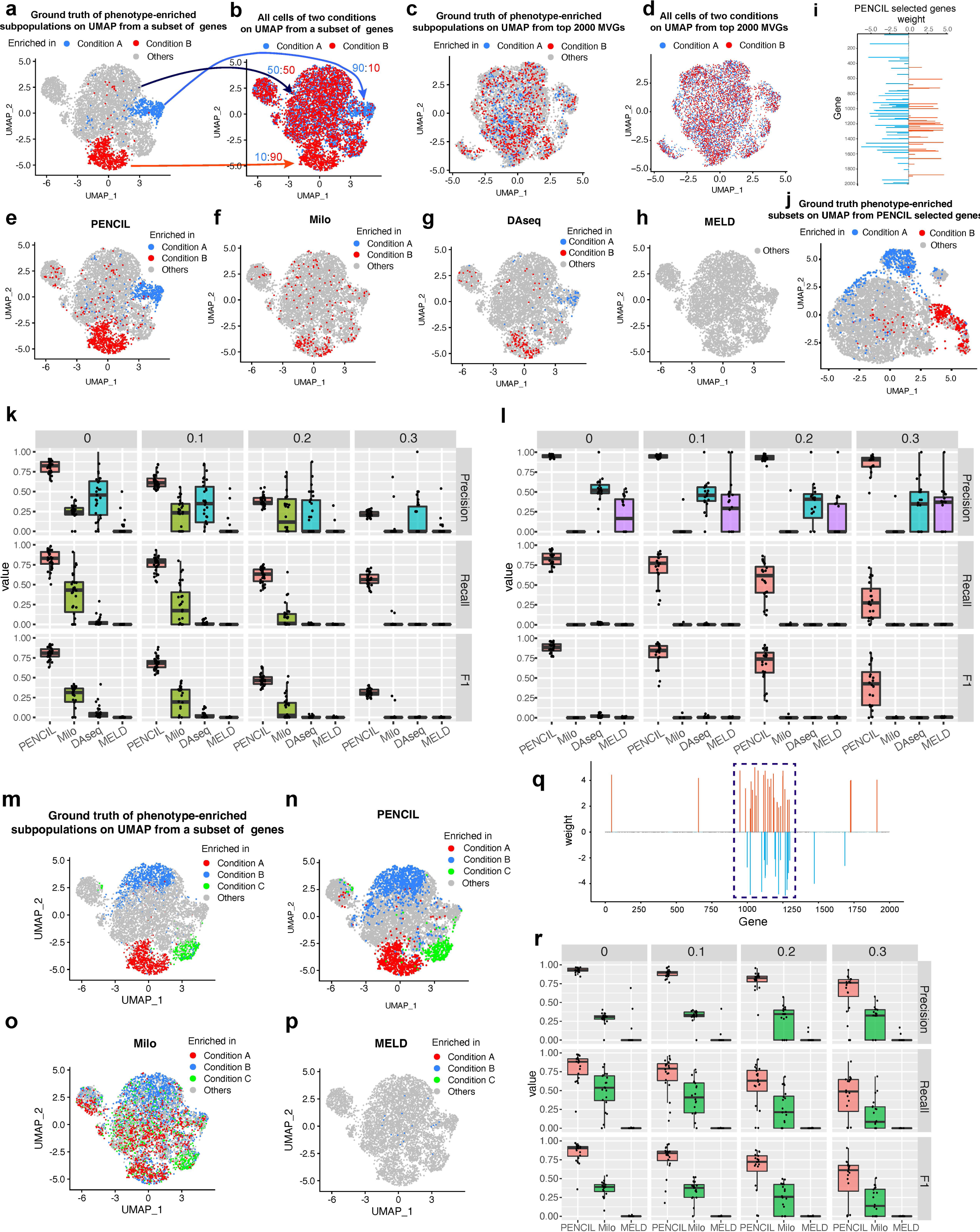
Evaluation of classification mode of PENCIL for simultaneously selecting genes and cells in simulations. **a**, The ground truth of phenotype-enriched subpopulations and background cells on UMAP generated from a manually pre-selected gene set (1000-1300th MVGs) for the simulation with two conditions. **b,** The two phenotypic subpopulations were assigned to the two conditions accordingly with a mixing rate of 0.1 and all other cells are evenly assigned with condition labels, as shown by the arrows and ratios. **c,** The ground truth phenotype-enriched subpopulations in panel **a** visualized on the UMAP using the top 2000 MVGs. **d,** The cells with condition labels in panel **b** visualized on the UMAP using top 2000 MVGs. **e-h,** The predicted results of PENCIL, Milo, DAseq and MELD. **i,** The learned gene weights by PENCIL. **j,** The ground truth of phenotype-enriched subpopulations in panel **a** visualized on the UMAP using the PENCIL selected genes. **k,** The box plots showing the comparison results of the four methods (*n*=30 simulations) with four different mixing rates 0, 0.1, 0.2 and 0.3. The evaluation metrics of precision, recall, and F1-score were calculated to assess the abilities to recover the simulated ground truth cell subpopulations. **l,** The box plots comparing the performances of PENCIL, Milo, DAseq and MELD in the simulated batch effects datasets with four different mixing rates (*n*=20 simulations). **m,** The ground truth of phenotype-enriched subpopulations and background cells on UMAP generated from a manually pre-selected gene set (1000-1300th MVGs) for the simulation with three conditions. **n, o, p,** The prediction results of PENCIL, Milo and MELD. **q,** The learned gene weights by PENCIL for the three conditions simulation. The dashed rectangle region indicating the pre-selected gene set (1000-1300 MVGs) to simulate the UMAP in panel **m. r,** The box plots of performance comparisons for PENCIL, Milo, and MELD in the simulations with three conditions and four different mixing rates 0, 0.1, 0.2 and 0.3 (*n*=20 simulations).

Due to its unique ability to simultaneously select genes and identify subpopulations, PENCIL recovered 84.5% of the ground truth phenotype-enriched cells while maintaining a high precision (0.833) (Fig. 2e, Supplementary Fig. 2a-c). In contrast, because the top 2000 MVGs were not able to capture the proper similarities of the ground truth phenotypic subpopulations (Fig. 2c,d), the other three KNN-based methods did poorly, especially MELD, which did not select any cells (Fig. 2f-h, Supplementary Fig. 2d). Indeed, the feature selection in PENCIL contributes to improving the performance of this process, as illustrated by the UMAP generated from the PENCIL selected genes, which captured an appropriate cell-cell similarity structure of the designed ground truth subpopulations (Fig. 2i,j). We repeated this experiment 30 times, each time with 300 randomly selected key genes from the top 2000 MVGs to cluster cells. Then, we picked out two clusters, designated them as two distinct ground truth subpopulations and placed other cells as background cells. We performed the label assignments for four mixing rates to mimic the varying components within subpopulations. We utilized precision, recall and F1 scores between the identified cells and ground truth cells to evaluate the four methods. As the mixing rate increased, the performances of all the methods decreased, but PENCIL consistently provided better performances than other methods (Fig. 2k). In addition, merging cells from different samples and conditions must address the batch-effect issue. Various batch effect removal algorithms have been developed to date^24^. PENCIL can take the batch-corrected and scaled expression matrix as input, such as the data processed by Seurat^25^. We exploited Splatter^26^ to simulate expression data with batch effect. The results suggested that PENCIL can be integrated successfully with classic batch correction methods implemented in the Seurat^25^ Package (Supplementary Fig. 3). We repeated the simulations 20 times with four mixing rates for the batch-effects and showed that PENCIL consistently performed better than existing KNN-based methods (Fig. 2l).

In addition, as noted before, PENCIL can naturally be extended to address multiple conditions. Therefore, we did similar evaluations on simulation datasets with three conditions (Fig. 2m, Supplementary Fig. 4a-c) using the same T-cell scRNA-seq dataset^6^ as the two conditions. For the comparisons, we included Milo and MELD because they can easily address more than two conditions, whereas DAseq can only handle two conditions.

Consistently, PENCIL outperformed other methods with 0.815 recall and 0.884 precision (Fig. 2n, Supplementary Fig. 4d,e), compared to 0.816, 0.001 (recall) and 0.418, 0.176 (precision) for Milo and MELD (Fig. 2o,p, Supplementary Fig. 4f,g), respectively. 80.4% of the PENCIL selected genes came from the manually pre-selected genes (1000th-1300th MVGs), which were used to generate this simulation (Fig. 2q), confirming its capability in feature selection to facilitate subpopulation identification. We repeated experiments in multiple conditions 20 times, demonstrating better performance for PENCIL than other methods (Fig. 2r).

Taken together, we evaluated PENCIL in identifying subpopulations of two conditions, three conditions, and datasets with batch effects. Given that our primary goal was to demonstrate PENCIL’s ability to solve the feature selection problem rather than claim superior performance to other methods, all simulations were designed to necessitate gene selection. In fact, when assessing performance based on a constant set of informative genes, e.g., genes learned by PENCIL, all methods performed comparably (Supplementary Fig. 5). Indeed, the feature selection function embedded in the PENCIL framework selected informative genes associated with phenotypes and helped improve the performance in identifying phenotype-enriched subpopulations hidden in a latent gene space, which cannot be accurately detected by methods lacking gene selection during the training process.

### PENCIL’s regression mode enables supervised phenotypic trajectory learning of cell subpopulations

In addition to categorical phenotypes, increasingly single-cell datasets are designed to profile tissues from multiple time points and continuous disease stages^27^, such as cell differentiation, disease progression and drug response^15–17^. Our LWR-based PENCIL framework can also easily incorporate those continuous phenotypes into the regression mode by updating the prediction loss function (Methods). In comparison to classic differential abundance analysis, which identifies the subpopulation enriched in each categorical condition only (Fig. 1k,l), regression-based PENCIL can reveal subpopulations undergoing a continuous transition between conditions (Fig. 1m). Herein, we conducted a series of simulations to demonstrate the performance and applications of PENCIL in the regression tasks. In the first simulation to demonstrate its utility, we used data from a real scRNA-seq T-cell dataset^10^ (16291 cells with 10 principal components) that had been processed by the principal component analysis (PCA) dimensionality reduction algorithm to generate time-point labels. Three overlapping time points on the selected cell trajectory were set as the ground truth for the simulation experiment (Fig. 3a, Supplementary Fig. 6a), and cell labels were simulated accordingly, with the other cells being randomly assigned a time label as background noise (Fig. 3b). Regressing the simulated time points as continuous variables, PENCIL captured practically the entire track of cells defined in the simulated ground truth (Fig. 3c, Supplementary Fig. 6b). Though Milo also claims to be able to handle continuous variables, it only picked out the cells at the beginning and end of the trajectory, omitting the middle cells (Fig. 3d). The Venn diagram comparisons showed that PENCIL did allocate more ground truth cells (92% vs 54%) with higher precision (90% vs 80%) than Milo (Fig. 3e). More importantly, the most unique characteristics of regression-based PENCIL is its ability to predict continuous time scores for the selected cells (Fig. 3f), whereas Milo merely tests for a decrease or increase (negative or positive) in abundance over time (Fig. 3g). The predicted continuous time orders of selected cells by PENCIL provide unique opportunities to make novel discoveries such as the gene expression pattern associated with the time orders. Intriguingly, in this example, the histogram plot of the distribution of the time orders predicted by PENCIL showed two additional peaks at time points 1.5 and 2.5, suggesting hidden cell transition stages between the 3 designed time points (t1.5, t2.5) (Fig. 3h). Thus, the predicted continuous time scores can reveal new critical time points or phenotypic stages between designated time points that would otherwise be either overlooked or unnoticed by experimental plans or clinical definitions.

**Fig. 3.**
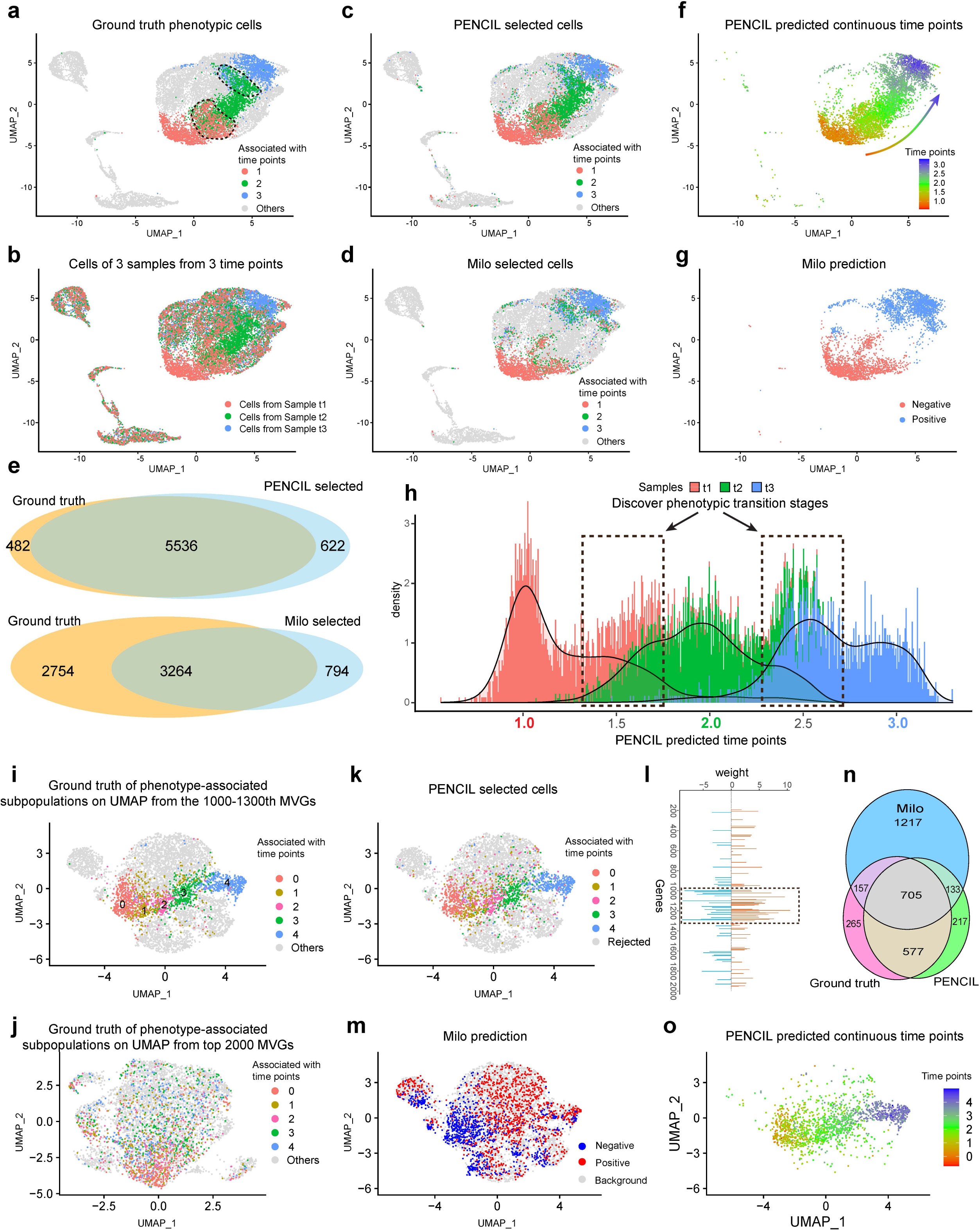
Evaluation of regression mode of PENCIL on the simulated datasets. **a**, For the first simulation, UMAP showing cells from a real scRNA-seq dataset assigned with 3 simulated ground truth phenotypic subpopulations and background cells. The regions within dashed lines indicating cells with labels evenly mixed by two adjacent time points. **b**, The 3 phenotypic subpopulations are assigned to the 3 samples accordingly and all other cells are evenly assigned to the 3 samples to form the sample labels for all cells. **c**, PENCIL selected cells. **d**, Milo selected cells. **e**, Venn diagrams comparing the cells selected by PENCIL and Milo with the ground truth phenotypic cells, respectively. **f**, PENCIL predicted continuous time points for the selected cells. **g**, Milo only assigned the selected cells as negatively and positively associated with the time course, corresponding to subpopulations decreasing and increasing with time, respectively. **h**, Histogram of PENCIL predicted time scores of selected cells colored by the sample labels. Dashed rectangles indicating the potential transition stages. **i**, For the second simulation, UMAP from a manually pre-selected gene set (1000-1300th MVGs) to show cells with simulated ground truth phenotypic subpopulations of 5 time points. **j**, Ground truth of phenotype-associated subpopulations in panel **i** visualized on the UMAP using top 2000 MVGs. **k**, PENCIL selected cells. **l**, PENCIL selected genes. The dashed rectangle region indicating the pre-selected gene set (1000-1300th MVGs) to set up the simulation in panel **i**. **m**, Milo predicted cells increase and decrease with the time course. **n**, Venn diagram comparing the cells selected by PENCIL and Milo with the ground truth phenotypic cells. **o**, The PENCIL-predicted continuous time points for the selected cells in the second simulation.

Next, we examined the gene selection function of PENCIL in the regression task. We employed the same scRNA-seq data of T cells^6^ in the classification tasks to simulate a time-series dataset. First, like in the previous experiment, we picked a subset of genes (the top 1000-1300th MVGs) from the top 2000 MVGs for the clustering and UMAP visualization to set up the simulated ground truth. Then we selected five subpopulations as the ground truth cells for five time-points and background cells based on the clusters generated from the selected genes (Fig. 3i). The standard top 2000 MVGs based analysis cannot correctly capture the structures of the five ground truth subpopulations (Fig. 3j). Then, we assigned the condition labels accordingly for phenotypic subpopulations and randomly assigned condition labels for background cells (Supplementary Fig. 6c). With the top 2000 MVGs expression matrix and the simulated labels as the input source data, the regression mode of PENCIL found most of the ground truth cells (Fig. 3k, Supplementary Fig. 6d) and the genes learned by PENCIL mainly located in the pre-defined 1000th-1300th MVG regions, as indicated by the dashed rectangle (Fig. 3l). In contrast, Milo selected many false positive cells (Fig. 3m). Specifically, PENCIL achieved 0.75 sensitivity and 0.79 precision, while Milo achieved 0.51 sensitivity and 0.39 precision (Fig. 3n). As before, the regression model of PENCIL can predict continuous time points for the selected cells to construct the trajectory (Fig. 3o). Additional simulations can be found in the accompanying supplementary material (Supplementary Fig. 6e-n).

By incorporating the supervised regression technique, PENCIL identifies high-confidence phenotype-associated subpopulations and orders them along a phenotypic trajectory, thereby facilitating novel insights into dynamic biological and pathological processes. Additionally, the gene selection function in PENCIL further empowers it to uncover continuous phenotypic patterns hidden within a latent gene space.

### PENCIL implementation, speed and scalability

PENCIL is implemented in Python to employ the powerful PyTorch framework enabling direct integration with other Python-based single-cell analysis platforms such as SCNAPY^28^. Alternatively, data preprocessed by R packages like Seurat can be saved into intermediate files for loading into Python. To streamline the analysis, we incorporated both native R and Python codes into a single document using "R Markdown", which allows us to seamlessly transfer objects between them. Thus, PENCIL can easily interact with Seurat^25^ and SCANPY^28^, two popular single-cell analysis frameworks. We provided tutorials to run PENCIL with SCANPY and Seurat. Furthermore, with the ever-increasing ability of single-cell sequencing to assess thousands to millions of cells^4, 29^, it is critical for the tool to analyze large-scale single-cell experiments efficiently. We simulated a large scRNA-seq dataset with 1,000,000 cells and 2000 genes from 3 conditions. We then down-sampled cells to run PENCIL in both regression and classification modes. The elapsed time, CPU and GPU memory usages increase linearly with the number of input cells to PENCIL (Fig. 4). When the full set of 1,000,000 cells were analyzed, the regression mode of PENCIL took less than 60 minutes, while the classification mode took 30 minutes. Both runtimes are acceptable for analyzing such a large dataset (Fig. 4a). As CPU and GPU memory were used to load data, regression and classification modes used the same amount for the same number of input cells (Fig. 4b,c). The runtime evaluations were performed using an AMD EPYC 7502 32-core processor and an NVIDIA A100 GPU.

**Fig. 4.**
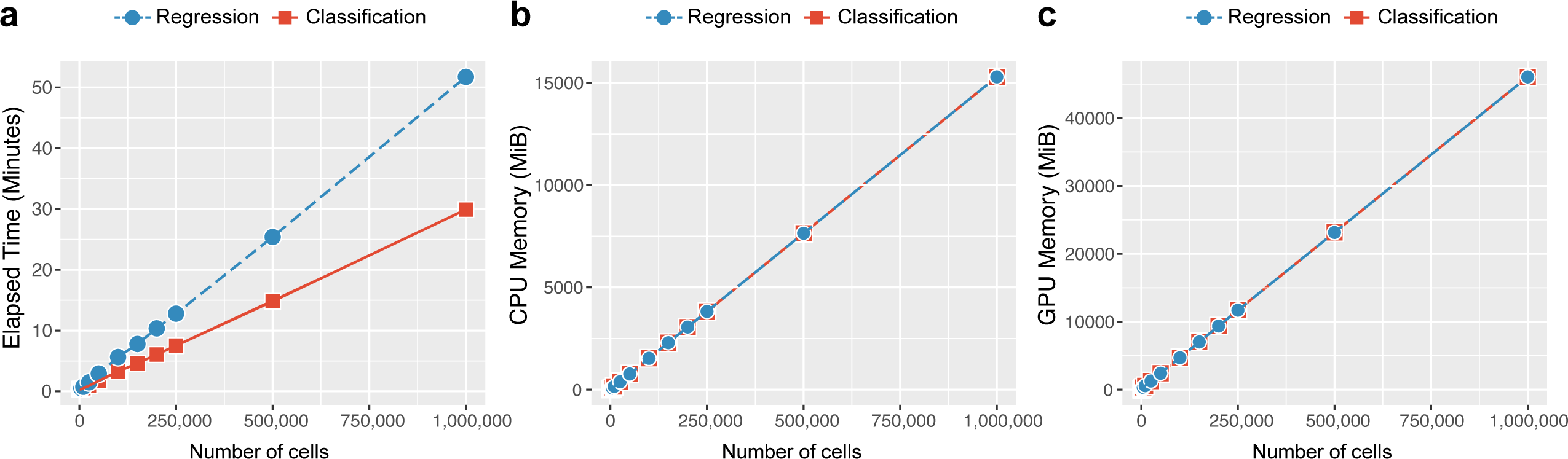
The running time and memory usages of PENCIL against the number of cells. **a**, Runtime of the PENCIL pipeline from inputting the normalized data to the final selected cells. **b-c,** Overall memory usage of CPU and GPU across the PENCIL workflow, respectively. MiB, mebibyte.

### PENCIL can identify T-cell subpopulations associated with immunotherapy outcome

To illustrate the utility of PENCIL outside of a simulated setting, we first applied PENCIL to a CD8 T-cell scRNA-seq dataset (6,350 cells) from melanoma patients consisting of 17 responders and 31 non-responders to Immune Checkpoint Blockade (ICB) therapy^6^ (Fig. 5a). ICB therapy has been a major breakthrough in cancer treatment^30^, but it only benefits a limited set of patients^31^. The purpose of this clinical dataset is to understand the underlying molecular mechanisms behind ICB response and resistance.

**Fig. 5.**
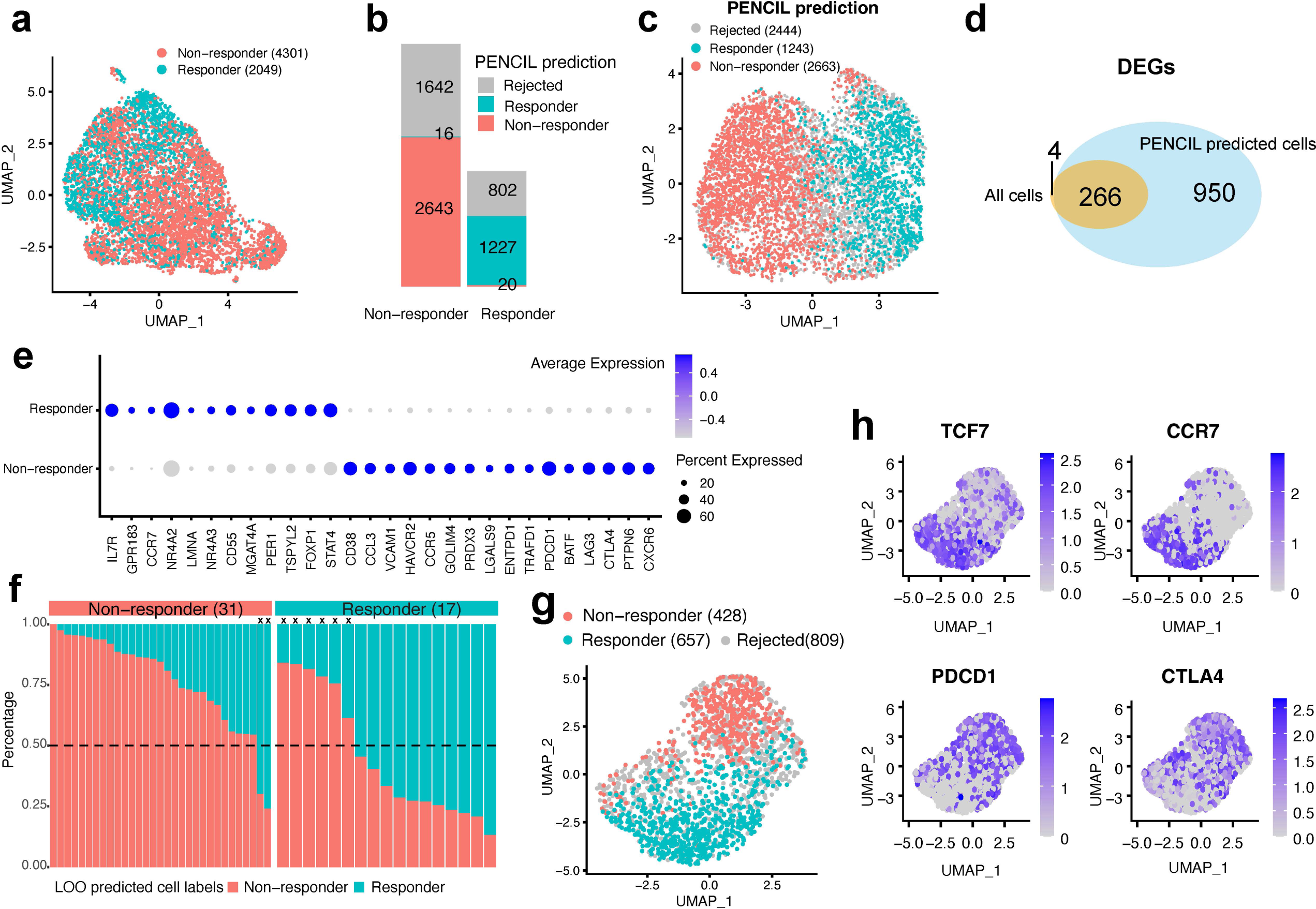
PENCIL analysis of T-cell subpopulations associated with melanoma immunotherapy outcomes. **a**, UMAP showing the cells using the top 2000 MVGs. Cell number in parentheses. **b**, The PENCIL predicted cell labels over the two conditions. **c**, PENCIL results on the UMAP based on PENCIL selected genes. Cell number in parentheses. **d**, Venn diagram comparing the DEGs of two conditions using all cells and the DEGs of PENCIL predicted labels of selected cells. **e**, Dot plots showing the expression levels of selected signature genes of PENCIL predicted phenotypes. The size of the dot encodes the percentage of cells expressing each gene and the color encodes the average expression level. **f**, Leave one out (LOO) prediction of responder and non-responder cells in the testing patient. The horizontal dashed line representing the cutoff to predict patients as responders or non-responders, and "x" indicating the LOO predictions inconsistent with the true condition. Sample number in parentheses. **g**, UMAP based on PENCIL selected genes during training showing the predicted labels of T-cells from a new melanoma patient in the Tirosh study^34^. Cell number in parentheses. **h**, The same UMAP from panel **g** colored by gene expressions of all T-cells from the Tirosh study.

Targeting the ICB outcome phenotypes, the classification mode of PENCIL identified 2,663 cells and 1,243 cells associated with the non-responders and responders, respectively (Fig. 5b). Simultaneously, PENCIL selected 88 informative genes (Supplementary Fig. 7), and the UMAP based on those selected genes exhibited a clear aggregation pattern for the PENCIL selected cells (Fig. 5c), showing how gene selection facilitated phenotypic subpopulation identification. To catalog transcription patterns underlying ICB outcomes, we executed a differentially expressed gene (DEG) analysis between the two subpopulations specific to ICB response and resistance. This analysis revealed 1,216 DEGs between the PENCIL selected phenotypic subpopulations (Fig. 5d), which included 950 new DEGs in addition to the ones derived from the original all responder vs. non-responder cells (Fig. 5d, Supplementary Table 1). Notably, the subpopulation associated with ICB responders has higher expressions of genes related to T-cell memory and survival, such as *IL7R*, *CCR7*, *LEF1*, *SELL* and *TCF7* (Fig. 5e). In contrast, the subpopulation associated with non-responders is marked by the expression of T-cell exhaustion and dysfunction genes such as *TOX*, *LAG3*, *ENTPD1*, *PDCD1*, *BATF* and *CTLA4*^32, 33^ (Fig, 5e).

Moreover, distinct from other strategies, our LWR-based supervised learning framework has an additional unique utility in that the trained PENCIL model from the given dataset can directly predict cell phenotypes from new single-cell samples, thus broadening the application of our framework. To demonstrate this utility, in the same dataset with 48 samples, we conducted a leave-one-out (LOO) evaluation of our PENCIL model. In this approach, 47 samples were used to train the PENCIL model, which was applied to predict cell phenotypes from the single left-out sample. We then classified each "left-out" patient as a responder if greater than 50% of cells were predicted as responder cells and evaluated this status against the actual clinical annotation. As a result, PENCIL correctly predicted the ICB outcomes in 40 out of 48 samples (Fig. 5f), which achieved 83.3% accuracy in the LOO evaluation, greater than 75% accuracy in the original study for the 48 samples^6^. In addition, given the PENCIL model trained on this T-cell melanoma ICB dataset, we applied it to an independent T-cell scRNA-seq dataset of a melanoma patient from Tirosh et al.^34^. In this new patient, PENCIL predicted more responder T-cells (657) than non-responder T-cells (428) (Fig. 5g), suggesting this melanoma patient would likely benefit from ICB treatment. The downstream marker gene analysis of the phenotypic subpopulations of this patient revealed that TCF7-high and CCR7-high Tumor-infiltrating leukocytes (TILs) were enriched in responder subpopulations while PDCD1-high and CTLA4-high TILs were enriched in non-responders (Fig. 5h). Thus, we demonstrated a unique function of PENCIL to transfer labels to new samples, which further independently confirmed the performance of PENCIL for phenotype-enriched subpopulation analysis.

### PENCIL learned the phenotypic trajectory of subpopulations in response to treatment

As previously discussed, PENCIL’s regression mode can resolve the phenotypic trajectory of subpopulations in a supervised manner that differs fundamentally from differential abundance analysis (Fig. 1 l,m). To illustrate this utility in real data, we next applied the regression-based PENCIL to a scRNA-seq dataset with samples collected at different times throughout a drug treatment period, which can provide insight into the mechanisms of action of a drug by characterizing transcriptional responses to the drug.

In a clinical trial to evaluate a NEDD8-activating enzyme (NAE) inhibitor in treating a mantle cell lymphoma (MCL) patient, a subtype of B-cell non-Hodgkin lymphoma (NHL), peripheral blood mononuclear cells (PBMCs) were collected from the patient at baseline and after 3 and 24 hours after drug infusion. Standard clustering of 3,236 PBMC cells detected 4 clusters with 3 B-cell clusters and one CD4 cell cluster (Supplementary Fig. 8a). The largest B-cell-1 cluster with 2,329 cells can be characterized by the deletions of chromosomes 6 and 9 through inferCNV^35^ analysis (Supplementary Fig. 8b), two recurrently affected genomic regions in MCLs^36^. Thus, we focused our analysis on this largest malignant B-cell cluster. In this cluster, standard clustering analysis based on the top 2000 MVGs did not find any cluster dominated by a specific time point (Fig. 6a, Supplementary Fig. 8c,d). We then performed PENCIL analysis by regressing the continuous cell labels 1, 2 and 3, corresponding to 0h, 3h, and 24h conditions, respectively. PENCIL identified high-confidence treatment-associated subpopulations, selecting 516 out of 1064 cells, 445 out of 583 cells, and 340 out of 682 cells from the 0h, 3h and 24h conditions, respectively (Fig. 6b). At the same time, PENCIL selected 44 informative genes (Supplementary Fig. 8e), and the UMAP plot based on this PENCIL selected genes clearly displayed the treatment response trajectory upon NAE inhibition (Fig. 6c,d). Then, correlating gene expressions with the predicted time orders of selected cells, we found 145 genes changing as cells progress along the treatment trajectory^18^ (Fig. 6e, Supplementary Table 2). Specifically, *JUNB* and *JUN*, whose overexpression is a hallmark of lymphoma cells^37^, had reduced expression following NAE inhibition (Fig. 6e). Overall, our PENCIL predicted time course analysis resulted in more signature genes than the differentially expressed genes (DEGs) of each time point from all cells (Fig. 6f). For example, gene *JUND* is positively correlated with malignant cell proliferation in NHL^38^, and PENCIL analysis found NAE inhibitor repressed its expression along the predicted time course during treatment (Fig. 6g), which was not detected by the DEG analysis (Supplementary Fig. 8f).

**Fig. 6.**
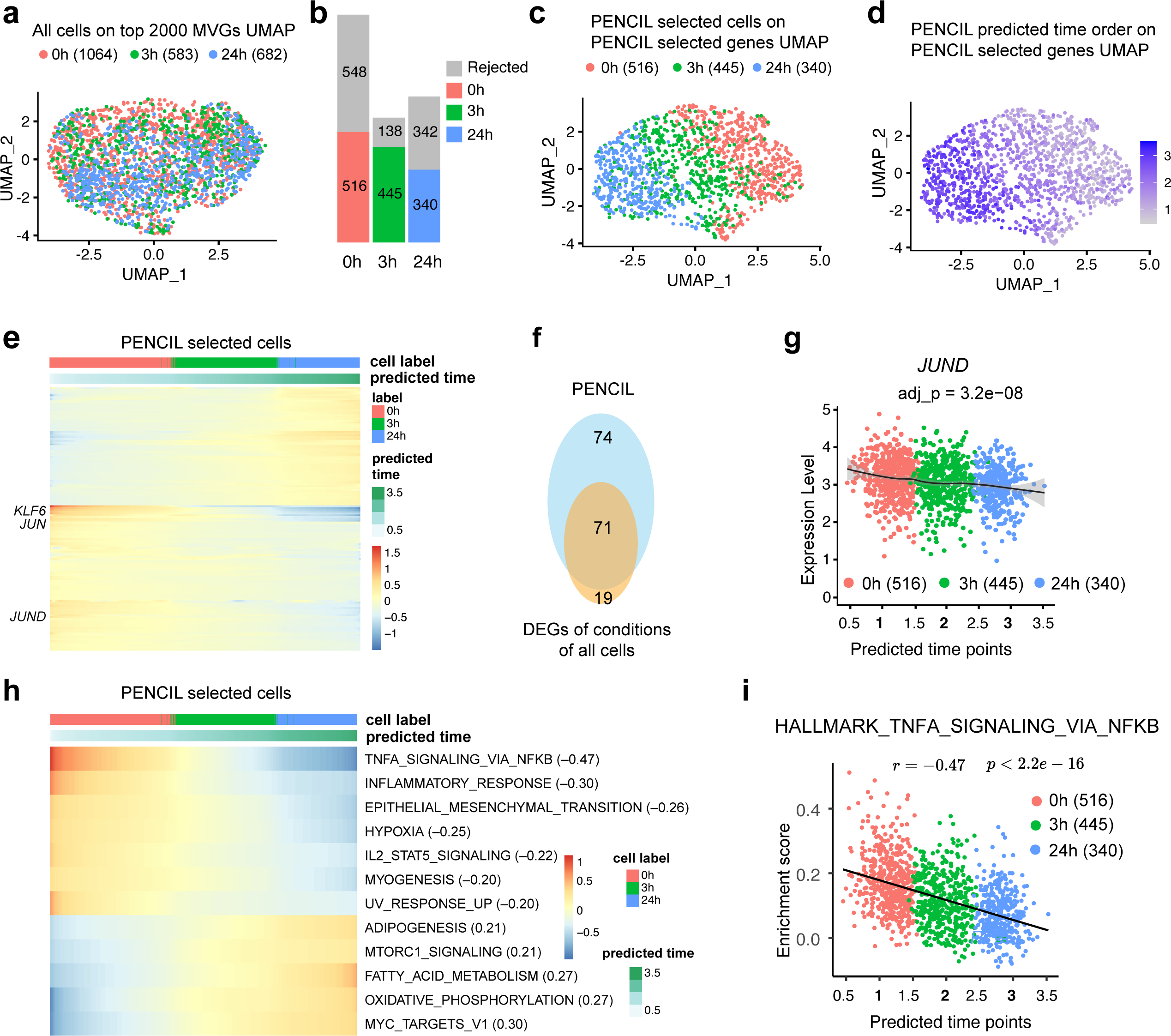
Regression mode of PENCIL analysis of scRNA-seq malignant B cells across 3 time points from an MCL patient. **a**, UMAP based on the top 2000 MVGs showing all cells of three conditions. cell number in parentheses. **b**, PENCIL selected cells across conditions. **c**, UMAP based on the PENCIL selected genes showing PENCIL selected cells colored by conditions. cell number in parentheses. **d**, PENCIL predicted time orders of PENCIL selected cells on the same UMAP in panel **c**. **e,** Genes significantly associated with the PENCIL predicted time points. **f**, Venn diagram comparing the DEGs of conditions using all cells and the genes associated with PENCIL predicted time orders. **g**, The scatter plot shows JUND as an example of genes significantly associated with predicted time points which were not detected by the DEG analysis. The adjusted P value was calculated by the Wald test. **h**, Hallmark pathways significantly associated with the predicted time orders with absolute correlation values great than 0.2. Pearson correlation values in parentheses. **i**, The scatterplot between the NFKB pathway activities and the predicted treatment time points predicted by PENCIL on cell subpopulations selected by PENCIL. The Pearson correlation coefficient and the corresponding P-value were indicated. The cell number is in parentheses.

Next, we explored the impacts of NAE inhibition at the pathway level. The proliferation and growth of MCL cells are dependent on NFKB signaling^39^. Interestingly, in our pathway analysis, the NFKB signaling pathway was the most negatively correlated with predicted time orders, suggesting NAE inhibition downregulated NFKB signaling along the trajectory to induce apoptosis in the MCL cells (Fig. 6 h,i). This observation is consistent with our pre-clinical data that NAE inhibitor abrogates NFKB pathway activity in chronic lymphocytic leukemia B cells^40^. Other on-target effects continuously downmodulated by NAE inhibition included the hypoxia pathway^41^ (Fig. 6h).

Together, this application demonstrated the unique abilities of PENCIL’s regression mode in selecting genes, selecting cells, and predicting time orders simultaneously, which unraveled the dynamic course of phenotypic changes.

## Discussion

PENCIL is unique in the following features and advantages (Supplementary Fig. 9). First, we introduced the learning with rejection strategy to single-cell analysis, enabling subpopulation identification in a supervised learning manner that is flexible to address categorical phenotypes or continuous variables. Second, we embedded the feature selection function into the supervised learning model, allowing for simultaneous gene selection and subpopulation identification to allocate phenotypic cell subsets hidden in a latent gene space that would otherwise be missed. Thus, we also introduced a new gene selection strategy to single-cell analysis beyond the existing unsupervised approaches. Third, the regression mode of PENCIL can select genes, identify phenotype-associated subpopulations and predict phenotypic trajectory simultaneously in a unified framework, providing supervised learning of subpopulations undergoing a continuous phenotypic transition. Fourth, by employing the powerful PyTorch framework, PENCI is fast and scalable, which can analyze 1 million cells within 1 hour (Fig. 4). Finally, besides subpopulation identifications, PENCIL has a unique utility that the model trained on the given dataset can directly predict cell phenotypes from new samples (Fig. 5).

The classification mode of PENCIL identifies subpopulations enriched by specific phenotypes, which has the same application as differential abundance testing algorithms like DAseq^10^, Milo^11^, and MELD^12^. However, our supervised learning-based PENCIL framework provides a more flexible way to select genes and identify subpopulations simultaneously from a global optimization perspective. To demonstrate this unique feature, the simulations for the comparison with other methods were designed in such a way that gene selection is necessary. However, we have to point out that our effort was not intended to develop a new method to improve the performance over existing methods incrementally, but to demonstrate that PENCIL is capable of performing gene selection to assist subpopulation identification.

Actually, when disabling the feature selection function, PENCIL and other methods performed similarly with the same input genes (Supplementary Fig. 5). Furthermore, the genes selected by PENCIL can be inputs for other methods to construct proper KNN graphs, which will be complementary to existing KNN-based approaches to improve their performances (Fig. 2f-h,o,p, Supplementary Fig. 5a,d) as well as utilize their advantages.

Although the extension of PENCIL to regression looks trivial, it has novel applications in single-cell analysis. Unlike the traditional supervised learning, in the LWR framework, this switch in loss function will affect not only the prediction term, but also the learning with rejection term, causing it to accept the cells transitioning between conditions (Fig. 1 l,m), which is a fundamentally independent application differing from differential abundance testing for single-cell data analysis. Thus, the regression mode of PENCIL extends beyond detecting static categorical cell states to reveal transitions during dynamic biological processes. Even though Milo can evaluate continuous inputs, it tends to select the subpopulations where phenotypic abundance monotonically increases or decreases, which usually misses phenotypic subpopulations in the middle of the time course (Fig. 3d,g). Most importantly, existing methods cannot assign time scores for the selected cells to reflect the dynamic course of phenotypes. Therefore, we believe the regression mode of PENCIL addresses a new application to supervised learning of the phenotypic trajectory of subpopulations.

PENCIL assigns cells from the same replicate with the same group label, so technical variability between samples is not taken into account, which is an inherited limitation in machine learning frameworks. In contrast, the statistics-based Milo can handle replication in an elegant way using the generalized linear model (GLM). Since PENCIL is complementary to other methods, we can provide the PENCIL-learned genes to Milo to exploit GLM’s statistical advantages. Furthermore, to address condition or sample imbalanced cell numbers, we introduced the condition/sample weights to the loss function, encouraging higher probabilities to retain cells from conditions/samples with smaller cell numbers.

As we stated before, our PENCIL framework is very flexible to take various forms of loss functions and we have implemented the loss functions to handle multi-category phenotypes and continuous phenotype scores. In the future, with single-cell experiments designed to profile more samples with survival information, we will add the cox-regression model into PENCIL to identify subpopulations associated with patient survival. Furthermore, though we only demonstrated the applications of PENCIL in scRNA-seq datasets, it can also handle other types of single-cell omics assays like single-cell ATAC-seq profiling different conditions^7, 42–44^.

In summary, by leveraging supervised LWR, we have developed PENCIL to simultaneously select genes, select cells, and predict categorical labels or continuous orders, thereby providing a new paradigm for identifying high-confidence phenotype-associated subpopulations from single-cell data. We anticipate that PENCIL will enable a broad application of phenotype-centric single-cell data analysis to deliver knowledge from single-cell experiments by focused interrogations of functionally and clinically significant cell subpopulations.

## Methods

### Learning phenotype-associated high confidence cell subpopulations by PENCIL

We build our method based on a concept known as Learning with Rejection (LWR), a machine learning strategy that introduces rejection labels in the prediction results (Fig. 1a,b). An insightful analysis for binary classification models with rejection was given in several previous studies^45–47^, and a general learning model with rejection has also been implemented experimentally^48^. For this application, we further develop a more robust and theoretically supported generic rejection-based learning method and apply it to single-cell data analysis to identify phenotype-associated subpopulations with high confidence. Moreover, we incorporate feature selection into this LWR framework to achieve the unique function of simultaneously selecting genes and detecting phenotype-associated subpopulations from single-cell data.

The workflow of PENCIL is represented in Figure 1c-g. The inputs for PENCIL are a quantified single-cell matrix and a label set of interest for each cell. Adhering to the general machine learning narrative conventions, let us denote the dataset combination to 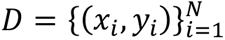, where *x*_i_ ∈ *R*^*d*^ is the *d*-dimensional gene expression vector of the *i*th cell and *y*_i_ is the corresponding target label of the *i*-th cell, such as condition, phenotype, stage, etc. (Fig. 1c).

Let *w* be a trainable weight vector on genes, *r*_Φ_ be a learnable model called rejector parametrized by Φ to determine the confidence score for the cells (*r*_Φ_(*x*) ≤ 0 means the cell has low confidence and it will be rejected, and conversely, it will be accepted), and ℎ_Θ_ denote the predictor to be learned with parameters set Θ (Fig. 1e,f). And *l* be the learning loss function for a specific supervised learning task. For any sample (*x*, *y*) in dataset *D*, PENCIL’s goal is to minimize the following rejection loss with gene weights (Fig. 1g),

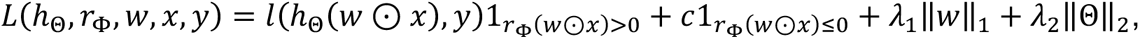

where ⊙ is the element-wise multiplication,1_*r*Φ>0_ and 1_*r*Φ≤0_ are indicator functions, and *c* is the cost of rejection. We impose a sparse penalty (*l*_1_-norm) on gene weights *w* to choose informative genes and *l*_2_-norm on Θ to control the model complexity of the predictor ℎ_Θ_, enable PENCIL to pick out high confidence cells that can be readily explained by a simple predictor.

The supervised loss *l* could come from a wide range of learning tasks, making PENCIL a flexible framework to be applicable in various scenarios. For example, if the target labels are multiple discrete categories, *l* can be a loss function for multi-classification; thus, PENCIL can identify the high confidence cell subpopulations related to multi conditions or phenotypes (Fig. 1j). When the labels are continuous variables, such as time points or disease stages, *l* can be a regression loss, so that PENCIL can determine a trajectory of selected cells highly correlated with the labels (Fig. 1m).

### Differentiable surrogate and model setup

The total loss function *L* cannot be optimized directly using the gradient-like algorithm, due to the inclusion of indicators 1_*r*Φ>0_ and 1_*r*Φ≤0_. We use (ℎ_Θ_) to denote (ℎ_Θ_(*w* ⊙ *x*), *y*) without causing ambiguity and temporarily ignoring the regularization terms. Drawing on the relaxation method in Cortes *et al.*^46^.

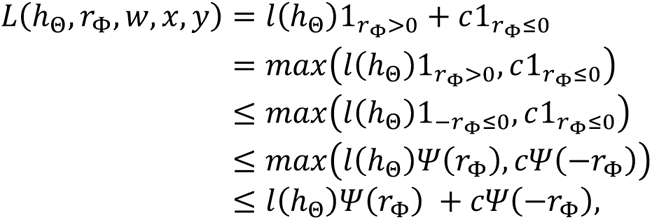

we can obtain the Max Surrogate (MS) and Plus Surrogate (PS) of *L* as,

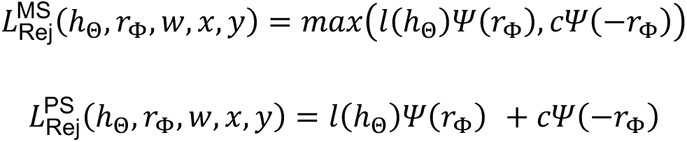

respectively, where ψ(·) can be any one of the forms mentioned in Charoenphakdee *et al*.^49^. Furthermore, the total loss on the whole dataset *D* can be formulated as

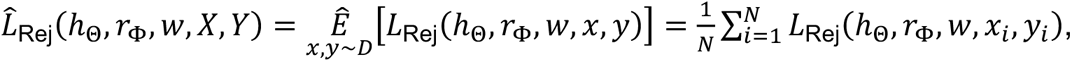

where *X* = (*x*_1_, … , *x*_*N*_), *Y* = (*y*_1_, … , *y*_*N*_), and 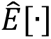 is the sample mean.

We substitute *w* ⊙ *x* with *x* in the latter part for narrative simplicity. In the context of a multi-classification (MC) task with *M* classes, the classifier ℎ_Θ_(*x*) is set to a linear classifier,

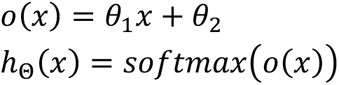

where *o*(*x*) ∈ *R*^*M*^. And (*x*) is a two-layer neural network using the activation function *σ*(*x*) = *x·tanh*(*softplus*(*x*))^50^, i.e.,

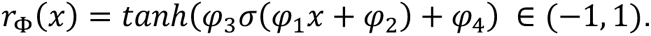

We use misclassification rate (MR) as the loss function for the multiclassification task, and set 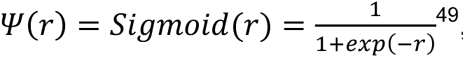^49^, and use PS type rejection. So, for multi-classification, our final implementation is

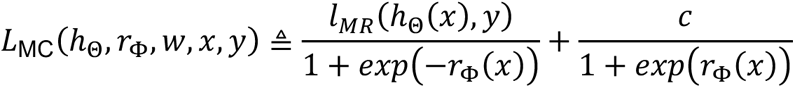

where *l*_*MR*_(ℎ_Θ_(*x*), *y*) = 1 − ℎ_Θ_(*x*)_*y*_, hence the selection range of *c* can be restricted to 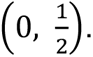.

Though binary-classification is a special case of multi-classification and is included in MR, we have also implemented some other losses dedicated to binary-classification, such as hinge loss^45, 48^.

In the regression (Reg) task, the regressor ℎ_Θ_(*x*) is set to a nonlinear neural network with a dropout layer,

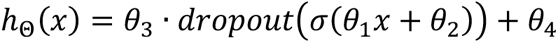

while ℎ_Θ_(*x*) ∈ *R*. The rejector *r*_Φ_(*x*) is the same as one in the classification task. The loss function for regression is Huber loss, ψ(*r*) = *Hinge*(*r*) = *max*(1 + *r*, 0)^49^, and MS type rejection is used, then,

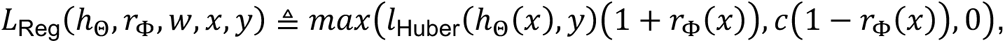

where

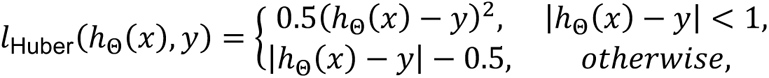

which is insensitive to outliers and gives more robust regression results than mean square error loss (MSE).

### Adjust cell numbers

In addition, we introduce class weights in the sample loss to overcome the class-imbalanced cell numbers, which is as follows,

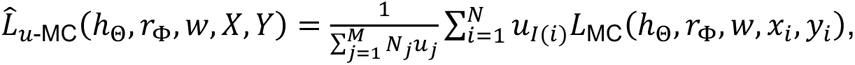

where *N*_j_is the number of cells in the *j*-th class, *u*_j_ is the weight for the *j*-th category and *I*(*i*) indicates the index of the category to which cell *i* belongs. Similarly, we can also define the weight of each sample to adjust sample-imbalanced cell numbers to have higher weights to keep the cells from samples with smaller cell numbers.

### Hyperparameter search

The rejection cost *c* is an important hyperparameter in the model. It directly affects the proportion of rejected cells and, hence, the final result. To eliminate the hassle of manual selection, we devised an algorithm to automatically select the hyperparameter *c*. The core principle is that when the labels are disrupted, the result of the rejection model should reject the vast majority of cells. Otherwise, it implies that the current cost of rejection is excessive, i.e., *c* is too large, hence a smaller *c* should be picked. On the other hand, to reject as few samples as possible on the original dataset, the rejection cost should be as high as possible. Thereby, we can take as the final choice the maximum cost that can reject the majority of samples on the dataset when the labels are disrupted. This search process can be accomplished by a bisection flow as shown in Alg. 1.

### Algorithm 1

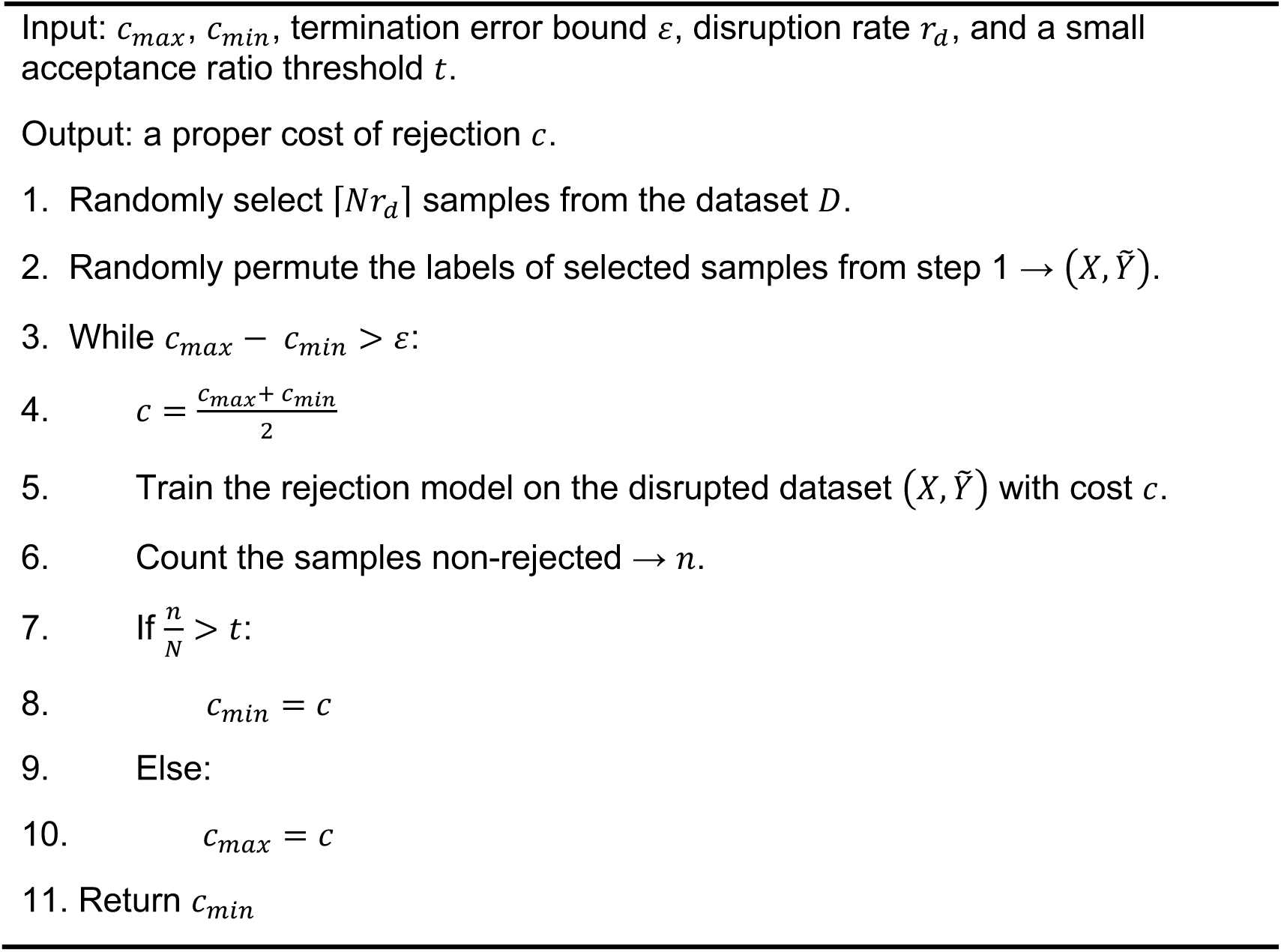

### Pre-train for faster convergence

The prediction model pre-trained on a purely learning task without the rejection module can converge faster in subsequent training. So, we first optimize (ℎ_Θ_) to pre-train the predictor ℎ_Θ_, and then optimize the rejection loss *ℒ* to train ℎ_Θ_(*x*) and *r*_Φ_(*x*).

### Simulation setup

In simulations for the classification mode of PENCIL, we exploited a real T cell scRNA-seq dataset^6^ with 6350 cells and 55737 genes. Since scRNA-seq data is noisy and sparse, we first selected the top 2000 most variable genes (MVGs) using the default function in the Seurat^25^ Package as the source data for PENCIL and other methods. First, for the specific simulations with two or three conditions as shown in Figure 2a,m, the 1000-1300th MVGs were manually pre-selected as the informative genes, then all cells were visualized and clustered based on the expressions of these pre-selected genes to generate the ground truth phenotypic subpopulations. After that, we picked out two or three clusters and designated them to be enriched in specific conditions, respectively. And all other cells were set as background cells. Next, we assigned simulated sample labels to the cells based on the conditions. We used a number *α* called mixing rate to control the ratio between the majority and the minority sample labels. Within each ground truth phenotypic condition, we assigned (1 − *α*) of the total cells of this condition with the designated majority condition labels, and the remaining cells with other labels. For the background cells, each cell was randomly assigned with a sample label. In this way, we got the labels for all cells for our analysis. We also depicted this simulation process in Supplementary Figure 1.

Second, to repeat simulations multiple times, we randomly selected 300 key genes from the top 1000 MVGs and subsequently clustered cells according to these pre-selected key genes. After that, we randomly picked out two or three clusters and designated them as the ground truth of phenotype-enriched subpopulations and placed other cells as background cells. Next, using the same procedure as before, we generated the condition labels for cells according to their designated ground truth phenotypes for four mixing rates (Fig. 2k).

For the simulation with batch-effects, we employed Splatter^26^ to simulate an expression matrix with 9000 cells and 8000 genes in two batches. 6000 of these cells are from one batch, and 3000 are from the other batch. And these cells are from 3 simulated groups with group probability of 0.6, 0.6, and 0.2. The probabilities of differential gene expression among the three groups were set as 0.1, 0.1, and 0.1. In order to produce the expression data which necessitates gene selection, we selected 500 genes and disrupted them 6 times along the cell orientation, resulting in 3,000 highly variable random noisy genes. Then, we merged these noisy genes with the original remaining 7500 genes into a new gene expression matrix of size 10500×9000. Following the default Seurat pipeline for finding MVGs^25^, we got the new top 3000 MVGs. As expected, most of these 3000 genes are the shuffled noisy genes, and only a very small fraction of them are key genes differentiating ground truth phenotype-associated subpopulations. Simulated groups can be completely separated under these differential genes (Supplementary Fig. 3a) and the batch-correction using Seurat revealed the 3 simulated groups (Supplementary Fig. 3b). But it did not work when using the top 3000 MVGs (Supplementary Fig. 3c). Thus, we obtained a simulated expression matrix comprising potential key genes, groups, and batches. Next, we generated the condition labels for all cells by setting the cells of group 1 as background cells, cells of group 2 and group 3 as two ground truth phenotypic conditions, and labeled them accordingly with a mixing rate of 0.1. After batch removal by Seurat^51^, using the batch-corrected and scaled expression matrix as an input, PENCIL selected the genes (Supplementary Fig. 3d) and identified 91.0% of the ground-truth cells with a precision 0.914, as shown in the UMAP generated from the PENCIL selected genes and Venn diagram (Supplementary Fig. 3e,f). To repeat this simulation, we conducted the simulations 20 times with 4 mixing rates and showed that PENCIL has better performance than other methods (Fig. 2l).

In the simulations for the regression mode of PENCIL analyses, we employed two types of single-cell expression data. In the first simulation, we used a scRNA-seq dataset preprocessed by PCA dimensional reduction^10^, which comprises 16291 cells and 10 principal components (PCs). Based on these principal components, we performed clustering and UMAP visualization following the standard pipeline in the Seurat^25^ package and selected 5 clusters (denoted as cluster1-5) as the ground truth trajectory (Supplementary Fig. 6a). We then set time-point labels for each of these selected clusters, where cluster 1,3, and 5 were assigned time point labels of t1, t2, and t3 respectively, while cluster 2 and 4 are set to be an equal mix of the two adjacent time point labels to mimic the transition stages (Fig. 3a). All of the other cells were set as the background cells, which were randomly assigned time point labels as noise (Fig. 3b). Then, we used the expression matrix with 10 PC along with the simulated time point labels to perform PENCIL analysis without the feature selection function. In the second simulation, because we wanted to demonstrate the feature selection of PENCIL in the regression mode, we employed the raw gene-level expression scRNA-seq matrix that was used in the classification tasks. We still pre-selected a subset of genes to necessitate the gene selection, which was further used for clustering and UMAP visualization to generate the ground truth subpopulations. For example, the top 1000th-1300th MVGs were used for clustering and UMAP visualization, which was used to select the clusters as ground truth subpopulations for the simulation case shown in Figure 3i. The time point labels of all cells were set up in a similar way as before by assigning time point labels according to their designated time point labels. To further demonstrate the regression mode of PENCIL’s capability in simultaneous feature selection, cell selection and continuous time points prediction, we performed two more simulation cases by manually pre-selecting different key genes (Supplementary Fig. 6e-n).

### Running Milo, DASeq and MELD

**Milo**^11^ samples a number of small clusters called neighborhoods from the KNN graph and then applies the negative binomial (NB) generalized linear model (GLM)^52^ to test differential abundance among conditions in each neighborhood. When using Milo, we set the neighborhood size parameter k to 30 and the sample probability to 0.1. Since Milo’s input must have multiple replicates to conduct statistical tests, cells from each condition were randomly divided into two replicates of equal size. We followed the tutorial of Milo to perform the analysis. Milo uses the spatially corrected false discovery rate (FDR) as the criterion to filter cell neighborhoods, and we set an FDR threshold of 0.05 to call neighborhoods that are differentially abundant between conditions.

**DAseq**^10^ is a multiscale approach based on the KNN graph to detect subpopulations of cells that are differentially abundant between single-cell data from two conditions. It calculates a differential abundance score vector for each cell based on the *k*-nearest neighbors of this cell across a range of *k* values, which is then utilized as the input to predict the biological condition of each cell using a logistic regression model. According to the tutorial offered by DAseq, we set the range of *k* to be 50∼500, with 50 as the step by default. DAseq subsequently picks the phenotype-enriched cells by setting a threshold on the score, which is derived by randomly permuting the labels.

**MELD**^12^ employs the theory of kernel density estimation on manifolds to compute the probability density distribution of biological states, which is then normalized to the relative likelihoods of the cells belonging to each state. The kernel density estimation method can also be viewed as a diffusion process of state labels on the graph. Then, the relative likelihoods are input into a Gaussian mixture model for cell clustering to identify phenotype-enriched cell clusters. Following the tutorial, we performed MELD analyses with default parameters for two conditions and multiple conditions.

### Evaluation metric: precision, recall, and F1 score

In all simulations, the ground truth benchmark is defined as the groups of cells that generate the phenotype-associated subpopulations. The true positive (TP) is the number of cells that are identified by both the evaluated methods and the ground truth cell set. The false positive (FP) is the number of cells selected by the methods but not included in the ground truth. The false negative (FN) is the number of cells rejected by the methods but belonging to the ground truth. Then, we use the precision, recall and F1 score to assess the performance of all methods, where precision is defined as TP/(TP+FP), recall is defined as TP/(TP+FN), and the F1 score is the harmonic mean of precision and recall, calculated by (2 * precision * recall)/(precision + recall).

### Standard scRNA-seq process in Seurat

We followed the standard Seurat (v4.0.5) pipeline to analyze scRNA-seq. After quality control and data normalization, the top 2000 most variable genes were selected by FindVariableFeatures function with default parameters in Seurat, which were further scaled. Then, principal component analysis (PCA) was applied to the selected MVGs to reduce noise from single-cell data for the downstream graph construction, clustering and low-dimensional visualization. The selection of the top most informative principal components was based on elbow and Jackstraw plots (usually 20-30). Data was visualized using the Uniform Manifold Approximation and Projection (UMAP)^22^ for dimension reduction, and clusters were detected by the FindClusters function with the default resolution (0.8). The differential gene expression analysis was performed for phenotype-associated subpopulations by the FindMarkers function in Seurat. Here, the default parameters for FindMarkers were Wilcoxon rank-sum test (two-sided), 0.25 for the log2 fold change cutoff, 0.10 for the parameter ’min.pct’, and adjusted p-value less than 0.05. When removing batch effects, we used Seurat comprehensive data integration pipeline^51^ to merge samples from different conditions.

### Sade-Feldman single-cell RNA-seq cohort with Melanoma immunotherapy outcome

Sade-Feldman cohort data of melanoma immunotherpay^6^ was used in this study. The gene expressions of single-cell RNA-seq were downloaded from GSE120575, consisting of 16291 cells and 36602 genes from 17 responders and 31 non-responders to Immune Checkpoint Blockade (ICB) therapy. The CD8+ T-cells (6350 cells) annotated in the original study^6^ were analyzed by PENCIL to identify high-confidence subpopulations associated with the ICB outcome (Fig. 5), which were normalized and scaled in the Seurat package. The scaled matrix of the top 2000 MVGs along with the ICB outcome labels were used as the input for PENCIL analysis. This CD8+ T-cell gene expression matrix was also employed to set up the simulation in different experiments (Fig. 2, Fig. 3i).

### Tirosh Melanoma single-cell RNA-seq data

The T-cell from Tirosh’s melanoma scRNA-seq data^34^ was predicted by PENCIL trained on another dataset to identify T cell subpopulations associated with immunotherapy outcomes. The preprocessed expression matrix was directly downloaded from GEO (accession number: GSE72056), and the 2,068 cells annotated as T-cells in the original paper were extracted for further analysis. Before performing the prediction, we excluded the smallest cluster with 174 cells characterized by the high expression of cell cycle-related genes, as indicated by another study that these cells may be contaminated with melanoma markers^6^. After that, we obtained 1,894 T cells for the final analysis (Fig. 5g,h).

### Genes significantly associated with predicted time points

We employed the functions implemented in Monocle3 (v1.2.9)^53^ to identify the genes significantly depending on the time points predicted by PENCIL’s regression mode. The gene expression levels were first fitted with the time points. Then, Wald test calculated the P-value by checking whether each coefficient differs significantly from zero, which was further adjusted by the Benjamini and Hochberg^53^. The genes were called as significant if their adjusted p-values were less than 0.05.

### Pathway analysis in single-cell RNA-seq

For each cell, we calculated the enrichment scores of the pathways in the MSigDB^54^ hallmark gene sets (v7.2) using the AddModuleScore function in the Seurat package^25^. Then, for each pathway, we calculated the Pearson correlation between the pathway enrichment scores and PENCIL predicted time points of PENCIL selected cells. The pathways significantly associated with the time course were called by absolute values of Pearson correlation coefficients greater than 0.2 and p-values less than 0.05.

### single-cell RNA-seq samples across three treatment time points from an MCL patient

This scRNA-seq dataset of an MCL patient across multiple treatment points was collected in a clinical trial led by Dr. Alexey Danilov to investigate the benefits of an NAE inhibitor^55^ on NHL patients. The manuscript of this clinical trial provides more detail about the protocol for generating scRNA-seq data, which is currently under review. We will upload this dataset to make it publicly available. In brief, we used the 10x Genomics Single Cell 3’ v3 kit according to the manufacturer’s instructions for the capture of single cells and preparation of cDNA libraries from patient peripheral blood mononuclear cells (PBMCs). The three samples collected at baseline and after 3 and 24 hours of treatment from the same patient were labeled with Cell Multiplexing Oligos (CMOs). Reads were de-multiplexed, aligned and counted using the 10x Genomics CellRanger v6.1.1 "multi" pipeline with default settings. After merging samples in Seurat, we performed data quality control by removing cell barcodes with < 200 UMIs, < 200 expressed genes or > 10% of reads mapping to mitochondrial RNA Genes. Doublets were removed using DoubletFinder^56^ (v2.0.3) with default parameters and a doublet rate threshold of 4%. We finally obtained the single-cell gene expression matrix with 14632 genes and 3236 cells. After normalization, the data was further scaled by regressing out the number of UMIs and the percentage of mitochondrial genes. The top 2000 most variable genes were identified with Seurat’s FindVariableFeatures using the default VST method, which were further analyzed by PCA. Then, the top 20 PCs were used to cluster and visualize the cells in UMAP. The cell types were annotated by SingleR^57^ (v1.8.1) following the standard procedure.

### InferCNV: Copy number alteration analysis from single-cell RNA-seq

InferCNV^35^ (v1.6.0) with the default parameters was used to predict the segmented copy-number alterations (CNAs) in scRNA-seq data. A healthy subject’s B-cells from the pbmc3k dataset in the SeuratData (0.2.1) were used as reference controls.

### Data availability

The description of public datasets used in this study and their accession numbers are detailed in the methods section above.

### Code availability

The open-source PENCIL program and its tutorials are freely available at GitHub https://github.com/cliffren/PENCIL.

## Acknowledgements

This work was supported by the following funding: the National Key Research and Development Program of China 2020YFA0712400 (to T.R. and L.Y.W.); NIH 1R21HL145426 (to Z.X.); Department of Defense Idea Development Award W81XWH2110539 (to Z.X.); Breast Cancer Research Foundation and NIH U01CA253472 and U01CA217842 (to G.B.M.); NIH 1R01CA244576 (to A.V.D.); NIH R35GM124704 (to A.C.A.); NIH R01CA250917 (to M.H.S.). We thank Dr. Jianming Zeng (University of Macau), and all the members of his bioinformatics team for generously sharing their experience and codes. We thank Weston Anderson for helping editing the manuscript.

## Author Contributions

Z.X. conceived the idea. T.R., L.Y.W. and Z.X. implemented the method and performed the analyses. T.R., C.C., A.V.D, X.G., S.D., L.Y.W. and Z.X. interpreted the results. X.W., M.H.S., A.C.A., P.T.S., L.M.C. and G.B.M. provided scientific insights on the applications. A.C.A. and G.B.M. contributed to the analytic strategies. L.Y.W. and Z.X. supervised the study. T.R., L.Y.W. and Z.X. wrote the manuscript with feedback from all other authors. All the authors read and approved the final manuscript.

