## Supplementary information for "Supervised learning of high-confidence phenotypic subpopulations from single-cell data"

[1] Academy of Mathematics and Systems Science, Chinese Academy of Sciences, Beijing, China. [2] School of Mathematical Sciences, University of Chinese Academy of Sciences, Beijing, China. [3] Computational Biology Program, Oregon Health & Science University, Portland, OR, USA. [4] Department of Biomedical Engineering, Oregon Health & Science University, Portland, OR, USA. [5] City of Hope National Medical Center, Duarte, CA, USA. [6]Department of Oncology Biomarker Development, Genentech Inc, South San Francisco, CA, USA. [7] Department of Cell, Developmental & Cancer Biology, Oregon Health & Science University, Portland, OR, USA. [8] Knight Cancer Institute, Oregon Health & Science University, Portland, OR, USA. [9] Department of Molecular and Medical Genetics, Oregon Health & Science University, Portland, OR, USA. [10] Division of Oncological Sciences Knight Cancer Institute, Oregon Health & Science University, Portland, OR, USA.

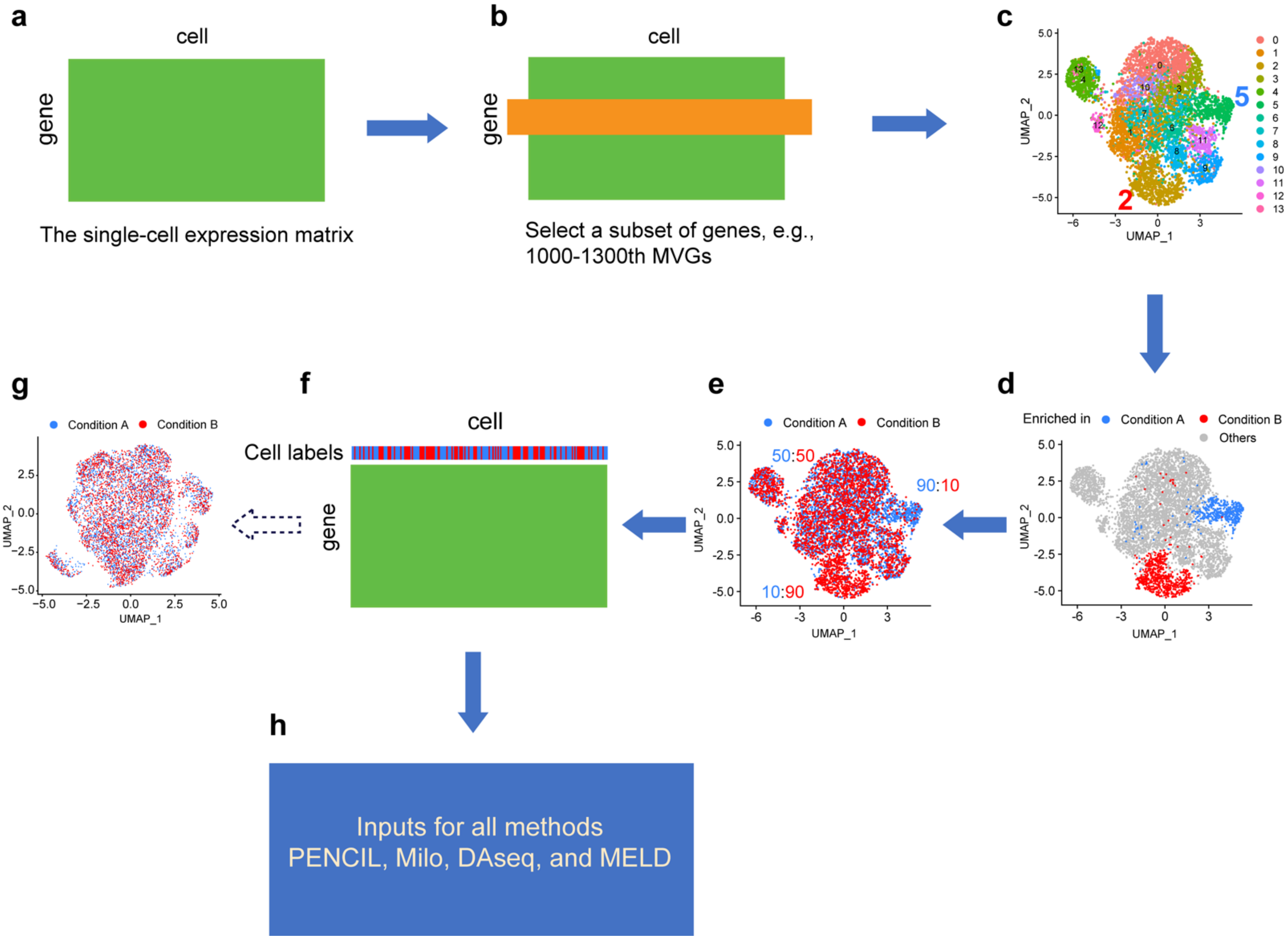


**Supplementary Fig 1. The simulation flowchart to necessitate gene selection. a**, The matrix from a real scRNA-seq dataset. **b,** Selecting a submatrix with a subset of genes indicated by the orange rectangle for the following clustering. **c**, UMAP visualization and standard clustering based on the submatrix from the previous step. **d**, Selecting two clusters (clusters 2 and 5) from panel **c** as the ground truth subpopulations enriched in the phenotypes, respectively. **e**, Assigning cells with condition labels based on the designed conditions in panel **d** and the given mixing rate. **f**, The raw matrix with each cell assigned with a condition label as indicated on the top bar. **g**, The UMAP using the top 2000 MVGs with cells colored by the condition labels. **h**, The raw expression matrix and cell condition labels as the inputs for all the methods.

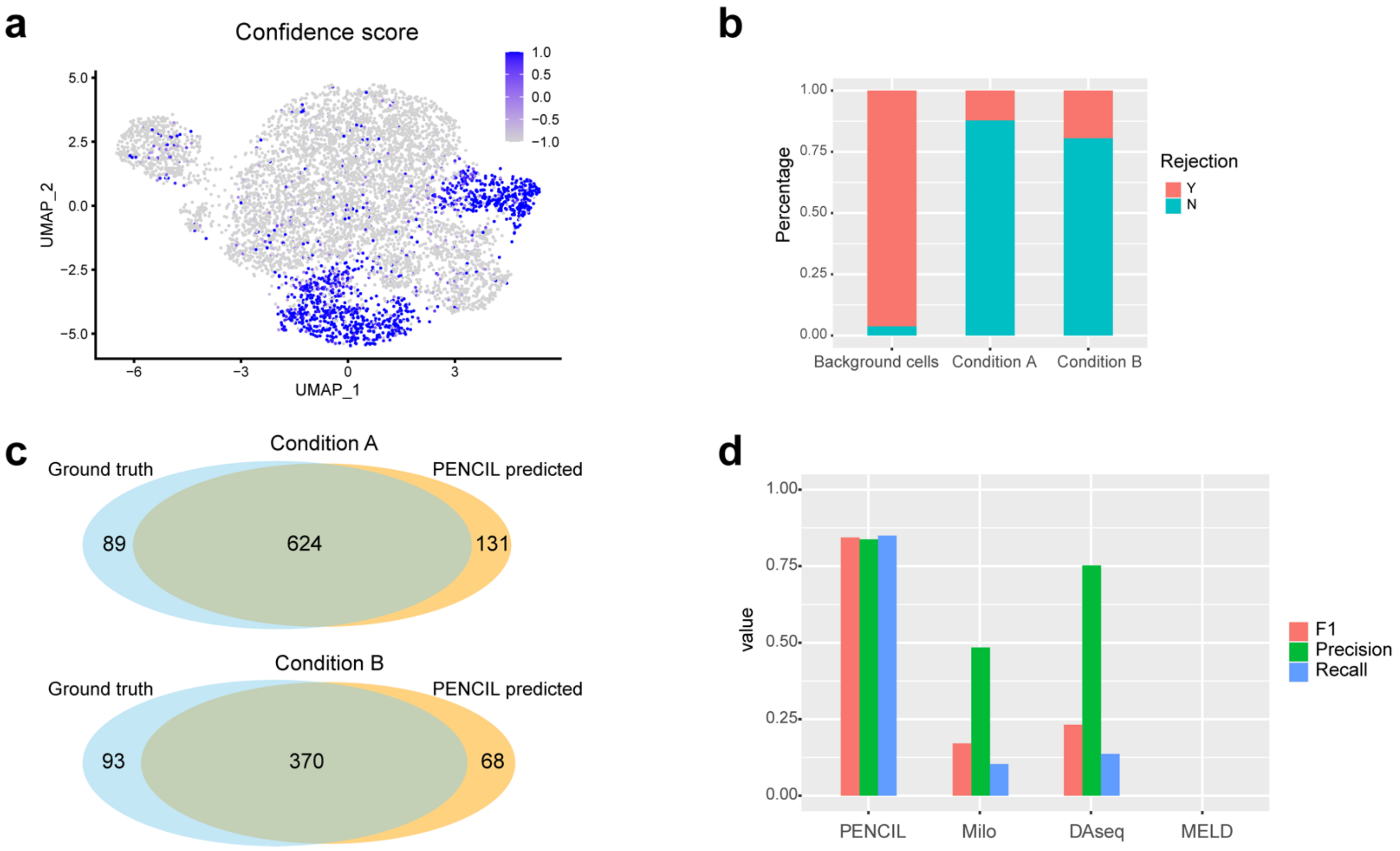


**Supplementary Fig 2. PENCIL classification analysis of simulated datasets with two conditions. a**, The confidence scores output by PENCIL. **b**, The distribution of the selected and rejected cells over the simulated ground truth of the conditions. **c**, The Venn diagram showing the overlap between the PENCIL selected cells and the ground truth phenotypic cells for the two conditions, respectively. **d**, The F1, precision and recall scores assessing the performance of the four methods.

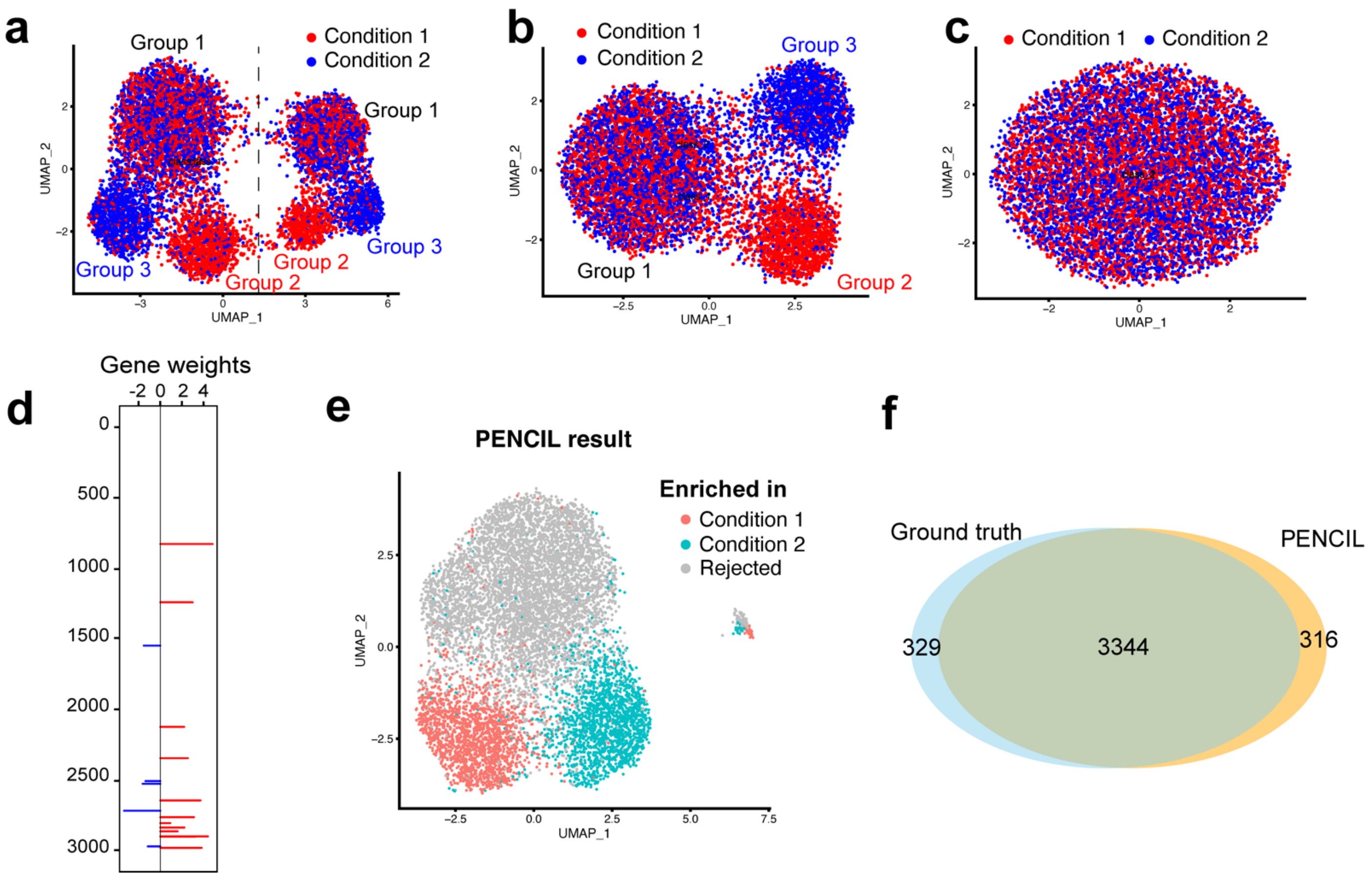


**Supplementary Fig 3. Evaluating PENCIL on the simulated dataset with batch-effect. a**, UMAP based on the manually curated genes showing the cells of two conditions from two batches separated by the dashed line. **b**, UMAP based on the manually curated genes showing the cells of two conditions after batch-corrections. **c**, UMAP based on the top 3000 MVGs showing all cells. **d**, PENCIL selected genes. **e**, UMAP based on the PENCIL selected genes showing the PENCIL selected cells. **f**, The Venn diagram showing the overlap between the ground truth phenotype-enriched subpopulations and the PENCIL selected cells.

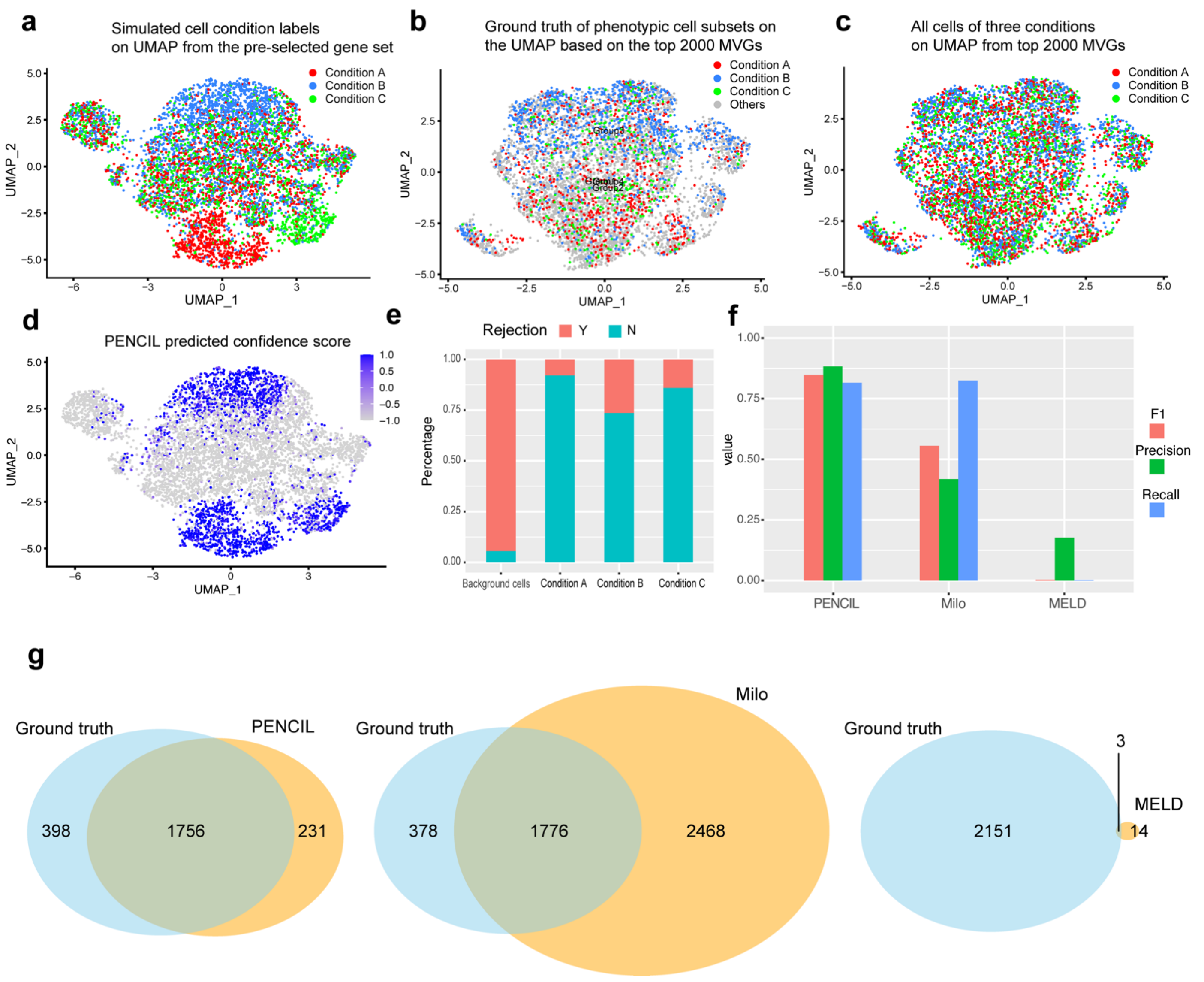


**Supplementary Fig 4. Evaluating PENCIL on the simulated dataset with three conditions. a**, The UMAP based on the manually pre-selected gene set colored by the cell condition labels generated from the ground truth cell subsets with a mixing rate 0.1. **b**, The ground truth phenotype-associated subpopulations visualized on the UMAP using the top 2000 MVGs. **c**, Cells with the same condition labels as the ones in the panel **a** visualized on the UMAP using the top 2000 MVGs. **d**, The UMAP based on the pre-selected genes colored by the PENCIL predicted confidence scores. **e**, The distribution of PENCIL selected cells over the ground truth cell conditions. **f**, The F1, precision and recall scores comparing the performance of the three methods on this simulated dataset with three conditions. **g**, The Venn diagrams depicting the overlap between ground truth cell subpopulations and cell subsets selected by the three methods, respectively.

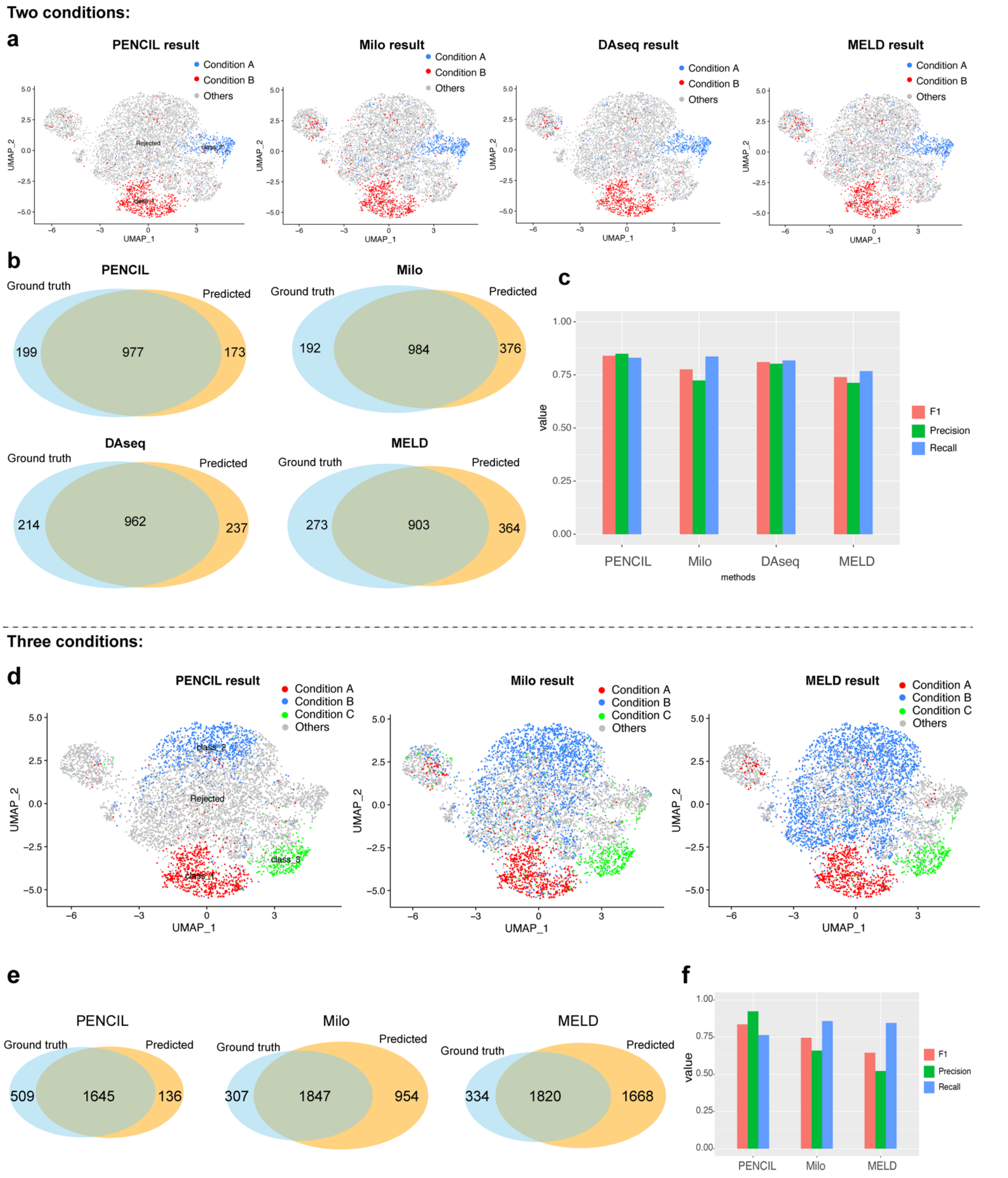


**Supplementary Fig. 5. Evaluating the four methods using the PENCIL selected genes as inputs. a**, The results of PENCIL, Milo, DAseq and MELD when inputting the genes selected by PENCIL for the two conditions simulation. **b**, The Venn diagrams comparing the result of each method with the ground truth phenotypic cell subpopulations. **c**, The F1, precision and recall scores assessing the performances of the four methods when inputting the genes selected by PENCIL. **d-f**, A simulation for the three conditions. **d**, The UMAP plots showing the results of PENCIL, Milo, and MELD when inputting the genes selected by PENCIL. **e**, The Venn diagrams comparing the result of each method with the ground truth phenotypic cell subpopulations. **f**, The F1, precision and recall scores assessing the performances of the three methods when inputting the genes selected by PENCIL in this simulated example with three conditions.

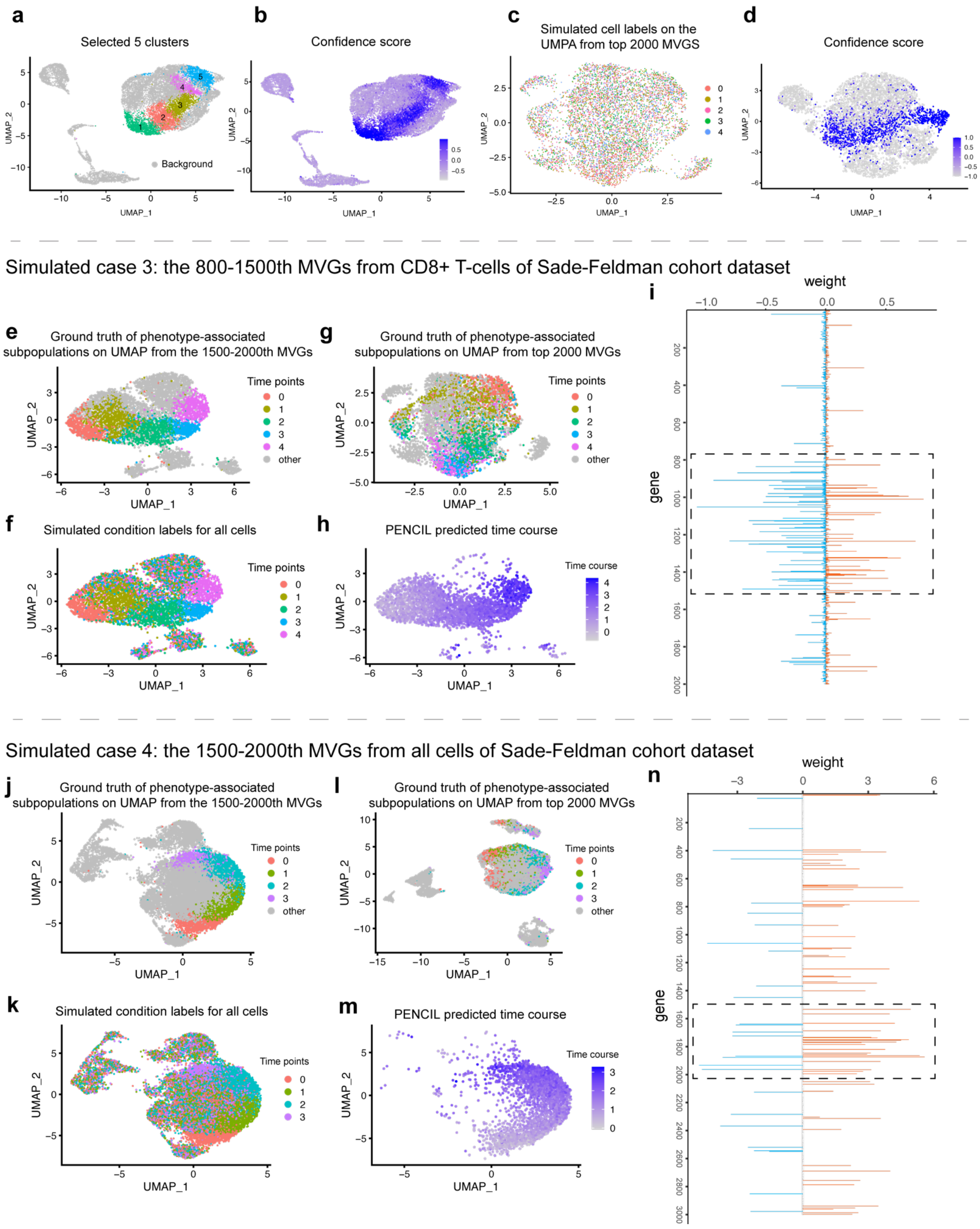


**Supplementary Fig. 6. Evaluating the regression model of PENCIL in simulated datasets. a**, UMAP showing the cells of 5 clusters selected as ground truth subpopulations corresponding to main Figure 3a. **b**, PENCIL predicted confidence scores on the UMAP corresponding to main Figure 3c. **c**, The cells with simulated condition labels visualized on the UMAP using the top 2000 MVGs. **d**, PENCIL predicted confidence scores on the UMAP corresponding to main Figure 3k. **e-i,** A simulated dataset for PENCIL regression analysis with manually pre-selected 800-1500th MVGs from the real Feldman T-cell dataset. **e**,

UMAP from a pre-selected gene set (800-1500th MVGs) to show cells colored by simulated ground truth phenotypic subpopulations of 5 time points. **f**, The 5 phenotypic subpopulations were assigned to the 5 samples accordingly and all other cells were evenly assigned to the 5 samples to form the sample labels for all cells. The UMAP is the same as the panel **e** colored by the cell condition labels. **g**, Ground truth of phenotype-associated subpopulations in panel **e** visualized on the UMAP using the top 2000 MVGs. **h**, PENCIL predicted continuous time points for the selected cells. **i**, PENCIL selected genes. The dashed rectangle region indicating the pre-selected 800-1500th MVGs for generating the UMAP in panels **e**, **f** and **h**. **j-n**, A simulated dataset for PENCIL regression analysis with manually pre-selected 1500-2000th MVGs from the Sade-Feldman cohort dataset. **j**, UMAP from a pre-selected gene set (1500-2000th MVGs) to show cells with simulated ground truth phenotypic subpopulations of 4 time points. **k**, The 4 phenotypic subpopulations were assigned to the 4 samples accordingly and all other cells were evenly assigned to the 4 samples to form the condition labels for all cells. The UMAP is the same as the panel **j** colored by the simulated condition labels. **l**, Ground truth of phenotype-associated subpopulations in panel **j** visualized on the UMAP using top 2000 MVGs. **m**, PENCIL predicted continuous time points for the selected cells. **n**, PENCIL selected genes. The dashed rectangle region indicating the pre-selected 1500-2000th MVGs for generating the UMAP in panels **j**, **k** and **m**.

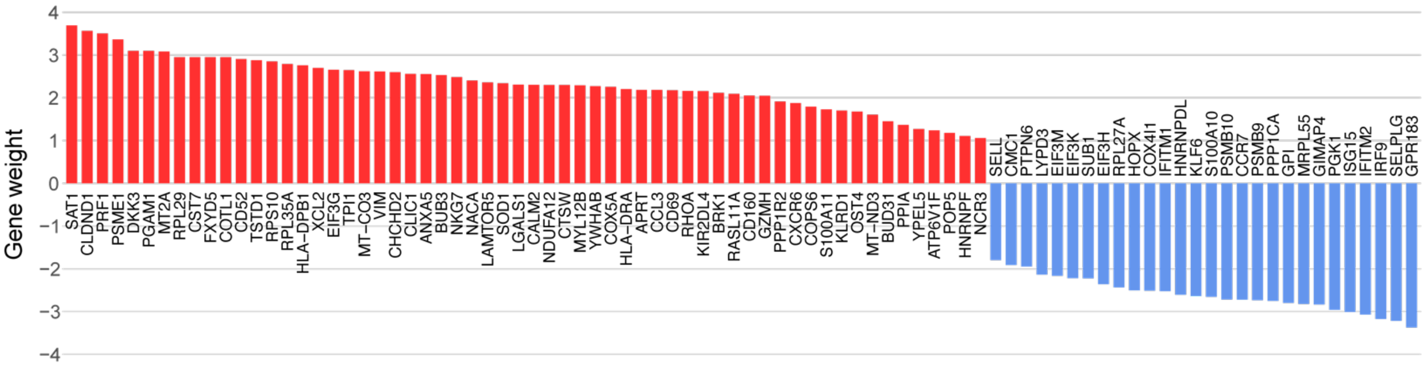


**Supplementary Fig. 7. PENCIL analysis of subpopulations associated with ICB outcomes.** The bar plot of estimated weights of PENCIL selected genes.

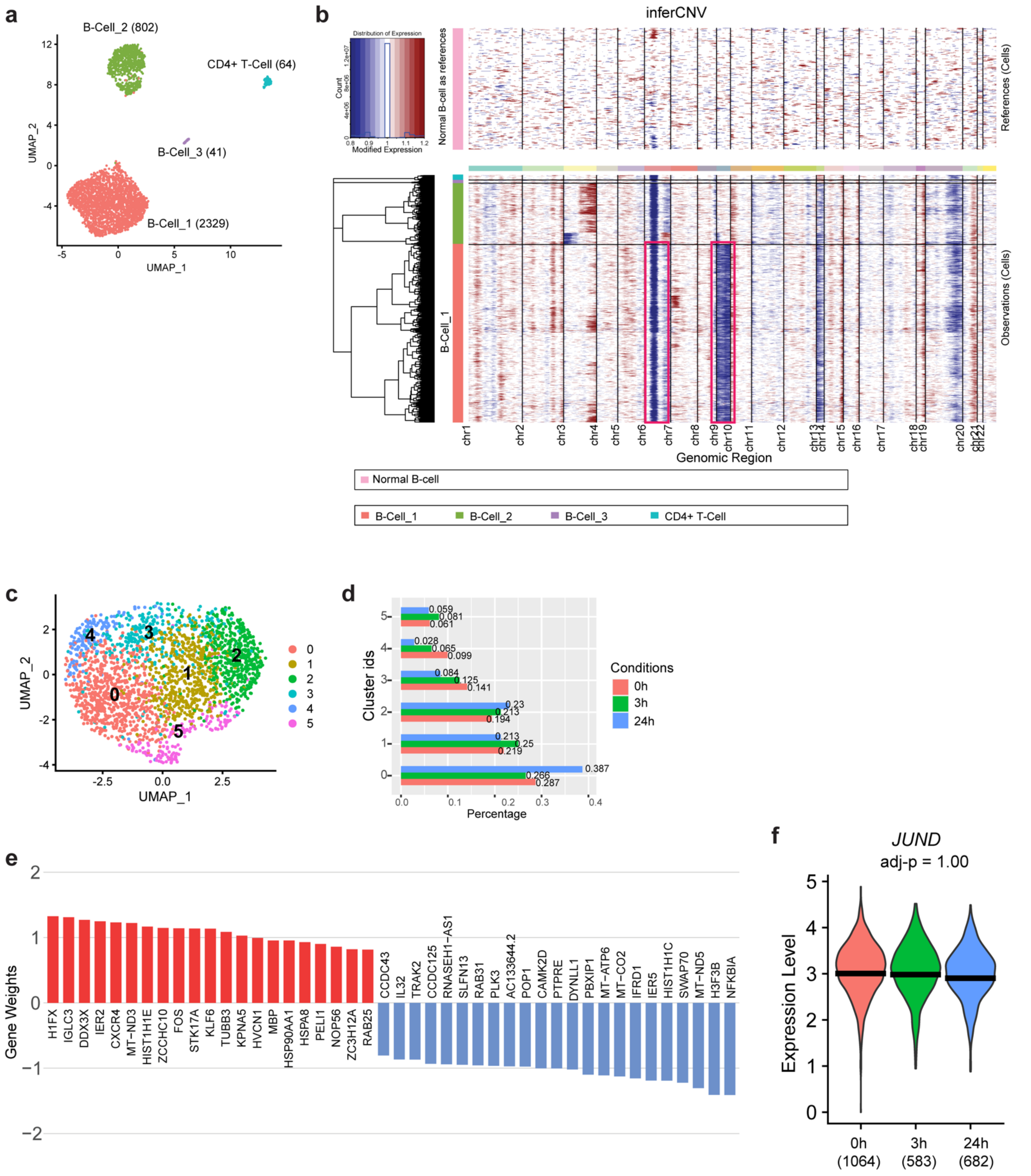


**Supplementary Fig. 8. Regression mode of PENCIL analysis of scRNA-seq malignant B cells across 3 time points from an MCL patient. a**, UMAP based on the top 2000 MVGs showing all cells of three conditions colored by immune cell types. cell number in parentheses. **b**, InferCNV analysis of B-cell clusters with normal PBMC B-cells as reference. Red rectangles highlight chr6 and chr9. **c-d**, Standard clustering of B-Cell_1 in panel **a** and percentages of conditions within each cluster. **e**, Bar plot of PENCIL selected genes with weights. **f,** Violin plots showing the expression levels of JUND on the original cells of three time points. The adjusted P value was calculated by the Wald test.

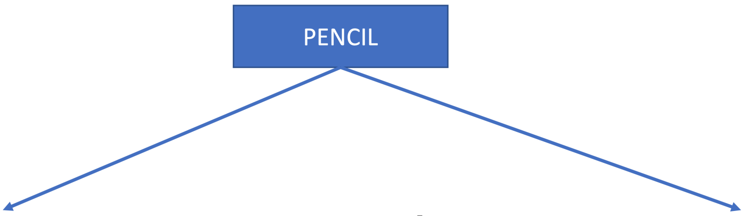


**a**

Classification
For an existing application:
Differential abundance (DA) of subpopulations

| Method | DA quantification | Clustering-free | KNN-free | Feature selection | GPU speedup | One million cells (in 1h) |
|---|---|---|---|---|---|---|
| PENCIL | Learning with rejection | ✓ | ✓ | ✓ | Fully | ✓ |
| Milo | Negative binomial GLM | ✓ | ✗ | ✗ | ✗ | ✗ |
| DAseq | Logistic classifier prediction | ✓ | ✗ | ✗ | Partially | ✗ |
| MELD | Kernel density estimation | ✓ | ✗ | ✗ | ✗ | ✗ |

**b**

Regression
For a **new** application:
Supervised phenotypic trajectory learning of subpopulations

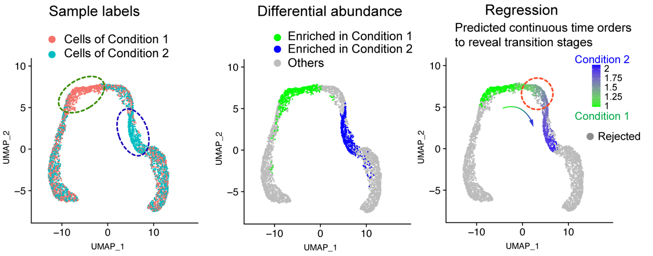


**Supplementary Fig. 9. A summary of PENCIL's two modes. a**, The advantages of classification-based PENCIL. **b**, The regression mode of PENCIL formulated a new application to reveal the continuous dynamic process of phenotypic subpopulations.

**Supplementary table legends**

**Supplementary Table 1.** The list of differentially expressed genes between PENCIL predicted cells associated with immunotherapy responders and cells predicted to be associated with non-responders. The p-values were calculated by Wilcoxon rank-sum test implemented in Seurat.

**Supplementary Table 2.** The list of the genes whose expression levels in the PENCIL selected cells significantly depend on the time points predicted by the regression-based PENCIL analysis on the MCL scRNA-seq dataset. The adjusted P values were inferred from the Wald test of the estimated coefficients as implemented in Monocle (see the Methods).
